# Non-decision time: the Higg’s boson of decision

**DOI:** 10.1101/2023.02.20.529290

**Authors:** A. Bompas, P. Sumner, C. Hedge

## Abstract

Generative models of decision now permeate all subfields of psychology, cognitive and clinical neuroscience. To successfully represent decision mechanisms, it is necessary to also assume the presence of delays for sensory and motor information to travel through the brain; but like the Higg’s boson in particle physics, directly observing this “non-decision time” from behaviour long appeared beyond reach. Here, we describe and apply a set of methods to empirically measure and characterise the properties of non-decision time in fast visually guided decisions (without requiring modelling assumptions). We gather 11 datasets from humans and monkeys from multiple labs and validate the method by showing that visual properties (brightness, colour, size) consistently affect empirically measured non-decision time, as predicted by neurophysiology. We then show that endogenous factors (pro-active slowing, attention) consistently do not affect non-decision time, in contrast to widespread reports based on model fits. Last, contrasting empirically observed non-decision time with estimates from the EZ, DDM and LBA models, we conclude that models cannot be generally trusted to provide valid estimates, either at a group level or for individual differences, and propose a hybrid approach that combines our empirical method with standard modelling.

---

The use of sensory information to guide action choices is a core principle of all mobile animals and a fundamental drive in evolution. Estimating how quickly and consistently information can travel to support this ability is both intrinsically interesting and essential to understand the neural underpinning of all sensorimotor decisions, in the healthy and diseased brains. In fast sensory-guided action decisions, most conceptual and modelling frameworks assume that choice and reaction time are co-determined in a competition between neuronal activity favouring different action options. This competition starts with the arrival of sensory signals and culminates with a point-of-no-return triggering one of the available motor responses. Before and after this process are inevitable delays, while sensory and motor information travel to and from the decision areas. Response times are therefore conceived to be the sum of (at least) three components: 1) sensory delay, 2) decision time and 3) motor output time (see Kelly et al., 2021; Servant et al., 2021; Weindel, 2021 for clear, recent articulations of this idea). As a result, the inferences drawn about decision depend inextricably on the assumptions or inferences made about the non-decision components (often treated together as ‘non-decision time’, NDT).

With little opportunity for directly measuring or validating the characteristics of NDT, nearly all the literature making use of decision models has relied on assumptions chosen for mathematical or computational commodity, and mainly validated by the model’s ability to capture a dataset, or through parameter recovery exercises (Ratcliff, 2013; Verdonck & Tuerlinckx, 2016). However, as we will delineate below, there are converging indications that what we have become accustomed to call ‘non-decision time’ in model frameworks does not capture the sensory and motor delay times as originally conceived. First, estimates vary greatly across studies, from null (which is biologically impossible, see for example Rohner & Lai, 2021) to above 1000 ms (which suggests the parameter includes more than basic sensory and motor delays, see for example Aschenbrenner et al., 2016; Powell et al., 2019). Another example relates to speed-accuracy trade-off, a long-standing decision concept in cognitive psychology. Humans and other intelligent animals strategically slow down when faced with an increased risk, or perceived risk, of making errors. Neuronal recordings in monkeys suggest that such strategic adjustments do not modulate the initial rise of visually-driven neural activity in the frontal eye field, one the key action planning areas (Heitz & Schall, 2012; Reppert et al., 2018). In contrast, decision modelling has repeatedly found strategic adjustments to affect the NDT parameter (see for example Dutilh et al., 2019; Lerche & Voss, 2018 and our discussion). If the NDT parameter truly captures non-decision time, this would indicate top-down influences all the way down to basic sensory processes or carrying on to motor execution, with fundamental consequences on our understanding of brain and cognition in general. However, if these indications imply that the NDT parameter captures processes other than non-decision time, this means that the models’ representations of decision is also flawed. To resolve these issues and questions, this article offers a route to an independent characterisation of sensory and motor time.

Besides its estimation being a necessary step to any confident understanding of decision processes, characterising non-decision time is also intrinsically valuable. In contrast to decision parameters (such as baseline, rise rate, threshold, or decision noise), *absolute* non-decision time is meaningful and interpretable in itself, irrespective of model architecture, and can therefore be easily compared across models, tasks, action modalities or species. Its components also have a straightforward relationship to direct (extracellular recordings) and indirect (EEG, MEG) measures of neuronal activity, and should relate to structural measures such as axonal diameter and myelination along specific tracts, or genetic factors affecting synaptic neurotransmission, providing a unique opportunity for critical external validation of these measures.

## Article Overview

Recently, we described a set of empirical methods to estimate the time taken for sensory information to travel from receptors to action decision brain areas (Bompas et al., 2020), and for motor commands to travel from the brain to the body (Bompas et al., 2017), all non-invasively, without relying on brain imaging or decision models. This line of research has so far exclusively focused on the dynamics of the decision competition, relying on small sample sizes with high number of trials, in conjunction with neural network modelling. Here we focus on the characteristics of non-decision time, and generalise this approach to multiple (mainly archival) datasets across different tasks and action modalities, and across humans and monkeys, providing benchmark expectations for NDT and solid foundations for sensorimotor research.

This work can form the basis of recommendations to decision model users regarding when, and how, NDT should be allowed to vary across trials and conditions, and when it may not, as well as what values should be considered plausible. Our main conclusions, applicable to fast visually guided decisions in response to clearly visible stimuli, are as follows:

- Sensory delay varies predictably with low-level properties of the signal such as brightness, colour and size, in line with neurophysiology (a conclusion we show to be inconsistently supported when using simple decision models).
- Sensory delay and output time are not affected by top-down adjustments such as slowing down due to different instructions or perceived error risk (at odds with the conclusions drawn from previous work using simple decision models, but in line with neurophysiology).
- In contrast to saccadic RT, output time for button presses is highly variable across trials and often positively skewed (at odds with the default assumptions in simple decision models).
- Empirically observed non-decision times show robust individual differences across conditions and tasks (that simple decision models struggle to capture on the same data).

Before we present the results underlying these conclusions, we use the rest of the introduction to review how NDT has been approached in the modelling literature and describe our empirical approach to estimating it.

## Non-decision Time in Decision Models

In the last decade, the use of simple decision models, such as the drift diffusion model (DDM, Ratcliff & Rouder, 1998) or the linear ballistic accumulator (LBA, Brown & Heathcote, 2008), has grown rapidly. Relatively easy to use, the parameters they provide attractively complement traditionally reported measures of mean RT and accuracy or choice, and the conceptual framework has helped explain – sometimes counterintuitive - patterns of behaviour across many types of task (Hedge et al., 2018). The goal of these models is to estimate, from the observed RT distributions of correct and incorrect responses, a small range of parameters that relate to the decision process. These parameters include a baseline level before the arrival of decision-relevant information, a mean rise rate capturing a sequential evidence-sampling process, a threshold capturing the amount of evidence sufficient to commit to a response, and a non-decision time as well as within and/or between-trial variability terms. All models rely on the assumptions that changes in the proportion of correct and incorrect RT responses and their respective latency distributions are diagnostic of changes within these underlying parameters.

Although very appealing, the success of this endeavour relies heavily on correct assumptions being made regarding the shape of trial-to-trial variability in the model parameters (Ratcliff, 2013). For instance, in the most prominent models – DDM or LBA – the starting point varies according to a uniform distribution, the rise rate follows a Gaussian distribution, while non-decision time is typically conceived either as a fixed value or a uniform distribution. As a result, the shape of most observed RT distributions can be captured by some combination of variability originating from these different sources. And by extension, such assumptions underlie the way in which differences between conditions, tasks or people are attributed to different parameters – decisional parameters or NDT, for example - which in turn generate theoretical conclusions about cognitive processes or the nature of individual differences or group differences (for example in caution/strategic control, information processing or simple sensorimotor delays). However, beyond their success at producing skewed unimodal RT distributions that fit observed ones, one needs to question the scientific grounds on which these premisses were built. This seems particularly important as the opportunity to externally validate those conclusions are rare. NDT is the one exception, where external validation is possible, at least for simple visuo-motor tasks. Before we explain this further, we will first review here how non-decision time has been handled so far from a modelling perspective.

Very few articles directly discuss the nature or properties of NDT. A search on PubMed for keywords related to the most common decision models (diffusion model and linear ballistic accumulator^1^) reveals above 11,000 entries since 1998, but only 1% of these articles make use of the words non-decision time (under various forms^2^) in their title or abstract (search completed on the 16^th^ of February 2023). Only a few of these studies are designed to specifically manipulate sensory and motor delays and assess the effect on model parameters. Manipulation of sensory conduction delay (Servant et al., 2016; Weindel, 2021), motor difficulty (Sandry & Ricker, 2022; Voss et al., 2004; Weindel, 2021) or response modality (Gomez et al., 2015) lead to the expected variations in NDT model estimates but, crucially, changes are also observed in the decision parameters. Reciprocally, when those *same* studies also manipulate factors expected to selectively modulate decision parameters (speed-accuracy trade-off or task difficulty), changes in NDT are also often reported. Overall, results such as these could either indicate an inherent permeability between sensory, decision and motor stages in the brain, or be diagnostic of poor specificity and imperfection in the model assumptions.

As for the studies where the focus was not directly on NDT, when changes in NDT are reported, they often mimic overall changes in mean RT, irrespective of whether these changes are theoretically expected or not. Most of the time, these changes occur alongside changes in other parameters. For instance, the well documented increase in RT with older age is partly accounted for by increased NDT (see Theisen et al., 2021 for a meta-analysis), and likewise for many other factors known to slow RT down, such as fatigue (Ulrichsen et al., 2020), sleep deprivation (Patanaik et al., 2014), alcohol (van Ravenzwaaij et al., 2012), dopaminergic drugs (Huang et al., 2015), task difficulty (Kelly et al., 2021), task switching (Imburgio & Orr, 2021), multi-tasking (Durst & Janczyk, 2019), and instructions favouring accuracy over speed (Lerche & Voss, 2018). Reciprocally, reductions in mean RT typical in Parkinson’s disease (Zhang et al., 2016), following practice (Dutilh et al., 2011), dopaminergic antagonist drugs consumption (Wagner et al., 2020), or speed instruction have all been associated with shorter NDT. Other conditions showing less systematic changes in reaction time, such as ADHD, also show less consistent changes in NDT, some reporting a decrease (see for instance Karalunas et al., 2014), an increase (Merkt et al., 2013) or no changes (e.g. Fosco et al., 2017; Karalunas et al., 2018). It is unclear to what extent these outcomes would be affected by different modelling assumptions and, as a general rule, the number of trials per participant or condition is too low to allow formal model comparison. Within this context, one may question the models’ ability to truly separate decisional and non-decisional contributions to overall RT changes.

This lack of conclusiveness echoes the excellent work by (Dutilh et al., 2019), who asked 17 research teams to use their favourite modelling approach to blindly identify the task manipulations modulating participants’ performance across 14 datasets. Despite none of these manipulations intending to induce a change in sensory delay or output time, changes in NDT were diagnosed in 37% of the inferences (89 out of 238). Even if researchers are not interested in NDT itself, not knowing when it should be constrained or not has consequences on other parameter estimates. For instance, it has recently been shown that estimates of between-condition differences in NDT and boundary separation (meant to capture caution) are negatively correlated (Grange & Schuch, 2021). This occurred in simulations when there were no real differences in either parameter. Trade-offs such as this will lead to incorrect conclusions about parameters judged to be of interest.

More generally, this under-specification reflects psychologists’ primary interest for understanding cognitive processes contributing to decision, rather than sensory delay or motor execution. When trial numbers are low, allocating several free parameters to capture non-decision time accurately could appear wasteful or even detrimental, as they will trade-off against free decision parameters (Boehm, 2018; Lerche & Voss, 2016). As a result, NDT in models is most often reduced to a fixed value (one free parameter) or a uniform distribution (two parameters, mean and range), when it is not simply ignored (i.e. set to zero following Carpenter & Williams, 1995; Smith, 1990). In contrast, output time is, like most biological processes, likely noisy with a positive skew when responses involve body muscles (button presses, pointing, reaching etc). Saccades are the only response where motor output time can be safely simplified to a fixed value (see Bompas et al., 2017, and Discussion). Previous parameter recovery work showed that ignoring skew can lead to systematic inaccuracies in decision parameter estimates (Verdonck & Tuerlinckx, 2016).

Despite raising awareness of this issue, possible consequences and solutions (Verdonck & Tuerlinckx, 2016; Weindel, 2021), modelling practices in the field are still commonly justified by claims that, in some contexts, assumptions surrounding NDT are of little consequence to the modelling outcome (Ratcliff, 2013). Parameter recovery estimates using the standard DDM showed that, when decision time is much more variable than NDT (e.g. four times), the distribution of their convolution hardly depends on the shape of the NDT distribution (Ratcliff, 2013; Ratcliff & Tuerlinckx, 2002). From this, many researchers may have concluded that all assumptions pertaining to NDT generally don’t matter. This overgeneralisation is unsatisfactory for several reasons. First, in many tasks, sensory and motor delays can contribute to a much higher proportion of the overall RT, either because the task is easy and leads to short decision times, or because the motor response required is longer and more variable. Secondly, it is unclear how conclusions based on parameter recovery exercises generalise to real data where effect sizes may be small and data quality low (small sample sizes, fewer trials or heterogenous populations as is common in clinical research). In real data, assumptions concerning NDT may well influence conclusions pertaining to other parameters, especially when these allow an effect to cross the traditional p = 0.05 threshold. Combined with selective reporting, modelling flexibility surrounding NDT parameters may lead to poor outcome validity. Third, previous simulation work focuses on the shape of NDT, but the plausibility of mean values and the factors that should be expected to affect or not affect NDT also need to be carefully considered. For instance, NDT estimates that are implausible (for instance close to zero or even negative, as observed in Rohner & Lai, 2021) should cast doubt on conclusions related to other parameters. Last, neglecting NDT appears less legitimate when alternative methods are available to constrain and validate NDT characteristics.

## Observing Non-decision Time Empirically

Sensory and motor delays can be estimated from visually triggered fast responses where one stimulus partially interferes with the initiation of an action to a previous stimulus (Bompas et al., 2020; Bompas & Sumner, 2011). The method typically involves two salient visual onsets separated by some delay, illustrated in Figure 1 using saccades (the same principles apply to manual tasks). Most often, the first signal is the target and the second one the interferer (e.g. a distractor or a stop-signal). Over many trials, the onset of the second stimulus creates a dip in the RT distribution to the first stimulus, when compared to the distribution in the absence of the interferer (black and grey curves on Figure 1B). The starting time of this dip (blue dot on Figure 1B) is precisely time-locked to the interferer onset (Bompas & Sumner, 2011; Reingold & Stampe, 2002). The time delay between the interferer onset and the dip onset indicates the non-decision time (we will refer to this measure as empirical NDT, or T_0_, Figure 1).

**Figure 1.**
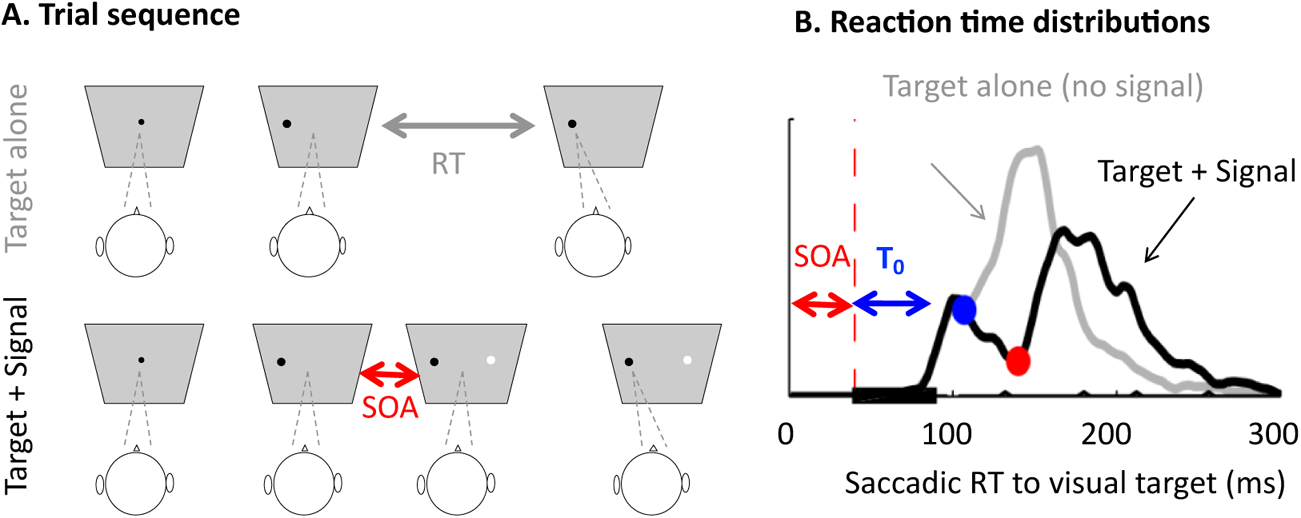
The visual interference method illustrated for an example study (from Bompas & Sumner 2011). **A.** The method compares the reaction time distribution to a target (here a peripheral black dot) in the absence (top row) and presence (bottom row) of a visual signal, here a task-irrelevant peripheral white distractor. **B.** Example RT distributions obtained from one participant in the target alone condition (grey) and when the signal onsets after the target (thick black curve), with an SOA of 40 ms and stays on for 50 ms. The thin black curve shows the very rare errors. The blue dot indicates the dip onset time, i.e. the time at which the signal-present RT distribution (black) starts diverging from the no-signal RT distribution (grey), as a result of the signal onset. The non-decision time (T_0_) is given by the time delay between the signal onset and the dip onset (70 ms for this participant).

The reason why the onset of the ‘dip’ (the transient interference in the RT distribution) reflects the non-decision time of the interferer stimulus is based on logic rather than mathematical assumptions. For a target signal to trigger an action, information about it needs to travel from the receptors to the decision area (sensory delay), rise to threshold (decision time), and translate into a movement (motor output time). In contrast, for an interferer to simply delay that action, it is sufficient that its signal reaches the decision area just before the *end* of the decision time, i.e. before the threshold is reached (Bompas et al., 2020; Salinas & Stanford, 2021). Any delay incurred by this interferer on decision time will only be measurable behaviourally after the output time has passed. Therefore, the earliest motor plan that a distractor will be able to disrupt is one whose RT is equal to the sum of the sensory delay of the distractor and the motor output time, i.e its non-decision time.

This logic does not need modelling *validation*, but it may please some readers to know that the only models able to convincingly reproduce the behavioural effects of visual interferers on RT distributions produce dips with onset perfectly matching the sum of the sensory delay and output time parameters. These models were exposed recently and do not constitute the topic of the present article. The interested reader may check Bompas et al. (2020) and Bompas et al. (2017) for more details. It is sufficient for now to emphasise that the key properties that allow these models to capture the interference effects are twofold: 1) they integrate time-varying exogenous and endogenous signals and 2) distant nodes mutually inhibit each other. While exogenous signals are conceived as automatic and mainly constrained by sensory events (stimulus onset and offset), endogenous signals are dependent on the task and instructions.

The saccadic modality is best suited for researchers interested in estimating sensory delay and investigating the factors that may affect it. Indeed, changes in empirical NDT for saccades can normally be attributed to sensory delay because saccadic output time shows negligible variance (see Discussion). This also means that saccade onset provides, on each trial, a nearly perfect read-out of the time each decision is reached in the brain. And once empirical NDT is known, it is straightforward to estimate the decision time from RT.

The interference method for manual responses typically supports similar conclusions as using saccades, but manual motor output time is longer and intrinsically noisy. T_0_ then indicates the shortest non-decision time in this noisy range. Researchers specifically interested in investigating motor output time distribution may use the method summarised in Figure 2, using saccadic and manual versions of the same RT task on the same participants. In this article, we provide the outcome of this process on a benchmark of 40 participants.

**Figure 2.**
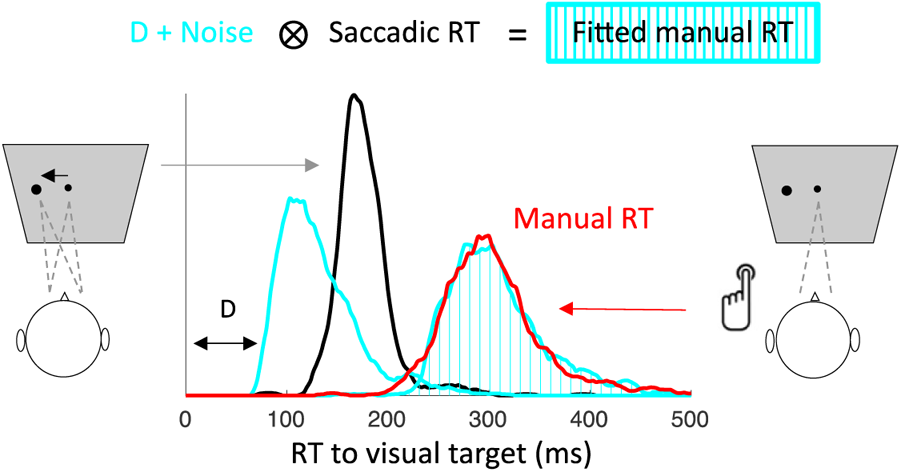
Estimating manual output time from RT on an example participant from Campbell et al. 2017. In simple detection tasks, manual RTs (red curve) are systematically longer and more variable than saccadic RTs (black curve). The manual output time distribution (cyan curve) is the distribution which, when convolved with the saccadic RT distribution, provides the best fit to the manual RT distribution. In this participant, the best-fitting distribution (cyan dashed area) is obtained using a gamma distribution with a shape of 2 (skewness of 1.4) and a scale of 28, added to a fixed delay (D) of 76 ms.

## Empirical Data—Methods

### Datasets Description

To illustrate which factors affect (or do not affect) non-decision time, the methods described above are applied to a range of empirical datasets. Datasets are chosen based on several criteria, rarely met together. First, they all require fast responses (saccadic or manual) to visual targets in the presence and absence of a second signal (e.g. distractor or stop signal). Second, they offer a within-subject comparison across at least two conditions leading to changes in response speed: different visual properties of the signal, different action modalities or different strategies (e.g. in datasets featuring the stop-task, trials following go-trials were compared to those following signal-trials, the latter displaying pro-active slowing, i.e. a change in strategy). Third, the number of trials per condition is sufficient (at least 200 trials per condition for distributional analyses and 100 for estimating the mean RT, typically more). Several such datasets were available internally, others were volunteered by other labs or requested from authors based on their publications. One new dataset was collected for the purpose of this article. A search on the OSF for additional relevant saccadic datasets was performed using the keyword combination ((“stop signal task” OR countermanding OR distract*) AND (saccad* OR “eye movement*” OR eye-movement*)) OR “saccadic inhibition”. It revealed 74 new projects, none of which matched our criteria. Datasets for human participants are presented below in chronological order, while the monkey dataset is presented first.

The **monkey** dataset covers a large part of the scientific careers of four rhesus macaques initialled A, C, Eu and X, trained to perform fast saccades to peripheral visual onsets, and stop on trials featuring a later visual onset (stop-signal). The behavioural and neuronal data from these monkeys contributed to many publications including Boucher, Palmeri et al. (2007) for monkeys A and C, and Sajad et al. (2019) for monkeys Eu and X, where interested readers can find a full description of the monkeys, task and data pre-processing. The total number of saccades made available to us were 5309, 4228, 12678 and 23298 for monkeys A, C, Eu and X. The ratio of signal-present trials was around 40% for A and C, and 50% for Eu and X. The SOAs varied across monkeys and covered a large range from 43 to 900 ms, with the number of trials per SOA varying across monkeys and recording sessions. Note that our analysis pipeline is largely immune to this variety (because it locks the response time on the onset of the interferer signal and pools across SOA), providing that a large enough number of trials are available at early SOAs where the early interference is visible, which was clearly the case here.

**Boucher et al. 2007a** (Boucher, Stuphorn, et al., 2007) had five participants take part in a saccadic, manual or bimodal visual stop-task. This dataset contributed to the post-signal slowing analyses in both modalities. Participants were asked to respond to peripheral visual onsets with either a fast saccade, a joystick move or both, and withhold their response when detecting a central visual onset. Signals were presented on 30% of trials, with SOAs 25, 75, 125, 175, 225, or 275 ms. For each participant and each modality considered, between 588 and 924 no-signal trials and between 252 and 396 signal-present trials were collected. The dataset also featured a selective stopping condition where participants needed to rely on the colour of the signal to identify which action modality should be withheld and which one was to be executed. This condition was not included in the present analysis, as there were not enough trials within each sub-condition to reliably extract dip onset.

**Bompas & Sumner (2009a)** compared saccadic behaviour to stimuli matched in salience but either visible or invisible to the magnocellular pathway (a manipulation known to slow sensory transmission time), with 5 participants. Stimuli consisted of achromatic luminance increments or chromatic shifts toward purple, amongst luminance noise. These stimuli were either used as saccade targets (first part of the experiment) or as distractors to be ignored (second part), where they were presented opposite to a black target. The first and second parts totalled 320 trials and 3240 trials per participant respectively. The SOAs used in the second part ranged from −80 to 80 ms in steps of 20 ms, but for the purpose of the present article, only positive SOAs were used to extract dip onset time. The number of useful trials per participant was then 450 distractor-present trials for each stimulus type (pooled across the 5 useful SOAs) and 2430 distractor-absent trials. The proportion of distractor-present was 76%. Data and code used in the present article are available [here](https://osf.io/ufja7/?view_only=abbccd30b6c24f8292c3b35783b65d25).

**Bompas & Sumner (2009b)** varied stimulus brightness with 3 participants. In a first part, participants performed simple saccades to light grey peripheral stimuli with 7 levels of luminance contrast (8 to 91%) against the darker grey background. 180 trials were collected per level and participant. In a second part, those stimuli were used as distractors and participants were instructed to ignore them while producing saccades to a black peripheral target randomly presented to the left or right of fixation. Distractors were always presented on the opposite side compared to the target, with SOA from −80 to 80 ms in steps of 20 ms and were present on 90% of trials. For the purpose of the present article, only positive SOAs were considered in order to extract dip onset. A total of 6300 saccades per observer were collected for part 2, including 450 trials for each level (pooled across the 5 useful SOAs) and 630 no-distractor trials. Data and code used in the present article are available [here](https://osf.io/rtnga/?view_only=cae9e376912e44ebade9823fd69de7f5).

The **free choice dataset** (unpublished) also varied stimulus brightness. A preliminary experiment was first conducted to select contrast levels for the main experiment. The design was the same as the first part in Bompas & Sumner (2009b) but with 8 brightness levels. In the main experiment, three levels were selected (the lowest, highest and one intermediary level from the preliminary experiment), and were presented as targets, either alone, or in pairs with varying SOAs from 0 to 60 ms in steps of 20 ms, where the participant was free to saccade to any of them under a speed instruction. The non-chosen stimulus, when appearing after the chosen stimulus, creates a dip in the RT distribution (in the same way as a distractor). The low and high brightness levels in the free choice dataset were the same as in the Bompas & Sumner (2009b) dataset, while the medium one was picked based on each individual’s RT, but was close to level 4 in Bompas & Sumner (2009b,0 and was therefore treated as such in the analyses. A total of 5300 saccades per observer were collected for part 2, including 900 one-stimulus trials at each level and 900 two-stimuli trials for each brightness pair. The data from the preliminary experiment were not used in the present article, as the latency to single trials during the main experiment was available and had more trials. Data and code used in the present article are available [here](https://osf.io/8d9je/?view_only=077cc915e3274c668f608632e178883c).

**Buonocore & McIntosh (2012)** manipulated distractor size and side (ipsi- or contralateral to the target, and therefore to the attended hemifield). Data from three of the four reported experiments are included here. Between 5 and 7 participants per experiment were asked to make fast saccadic responses to a right peripheral onset, in the presence or absence of a distractor stimulus, meant to be ignored. Only one SOA was used per participant, ranging from 69 to 234 ms, which corresponded to the individual’s median RT during a preliminary phase minus 90 or 110 ms. Each condition was sampled with between 150 and 400 trials per participant. Experiment 1 and 2 used only contralateral and ipsilateral distractors respectively, with varying sizes. Experiment 3 used both contra and ipsilateral distractors over a smaller range of sizes. Experiment 4 was ignored here, as it included only one distractor size and target side was random, and therefore did not meet criterion 2 for our analyses (having conditions with different mean RT, or otherwise known to affect processing time).

**Gulberti et al. (2014)** had 6 participants take part in a saccadic, hand pointing or bimodal stop-task, while electromyograms (EMG) activity was recorded. In separate blocks, participants were asked to quickly saccade to a visual peripheral onset (while keeping their hands still) or point with their finger (while keeping the eyes fixated) or do both. On 25% of trials, an auditory stop-signal was presented, with the instruction to withhold any response. Because of the auditory nature of the signal, we only used this dataset to investigate the motor execution time (delay between the first detectable muscle activity from the EMG and RT), based on 1800 no-signal trials per participant per condition (unimodal and bimodal).

**Campbell et al. (2017)** had 40 participants take part in a saccadic and manual version of the same task, before and after both alcohol and placebo drink conditions, and under instructions to either ignore or stop to a visual signal. All the analyses presented here were made after pooling across the three sober blocks (pre-placebo, post-placebo and pre-alcohol). This dataset was used to assess the effect of instruction on NDT in both modalities and characterise individual differences in NDT and manual output time. Participants were instructed to respond to a white peripheral onset with either a saccade or a left/right button press. On 25% of trials, a red central stimulus appeared after the target, with SOA 50, 100, 167, 233, 317 or 400 ms. In some blocks participants received the instruction to ignore the signal while, in other blocks, their task was to withhold their response. For each combination of action modality, instruction, and drink, 216 signal-present trials were collected across the six SOAs, and 648 no-signal trials. Although participants were instructed not to wait for the signal in the stop blocks, the contrast between the two blocks revealed clear pro-active slowing. We used the data available on the repository indicated in the original article, which only included RT for correct responses. Data and code used in the present article are available [here](https://osf.io/e5shq/?view_only=a788101082ff4b208ffc40e3b091b94f).

The experiment 2 in **Bompas et al. (2017)** had 4 participants doing a saccadic and manual version of the same task. Participants were instructed to respond to a peripheral black target onset randomly presented to the left or right, either with a saccade or a button press, in separate blocks, as fast as they could. On 83% of trials, a white distractor would appear at the opposite location from the target and participants were instructed to ignore it. Only the no-signal trials were considered here and were included in the manual output time analysis (as there wasn’t another experimental manipulation affecting RT). The number of trials collected per condition were 500. Data and code used in the original and the present articles are available [here](https://osf.io/vh8wy/?view_only=7a6d1fa67eb542eca3e96251fbafe4f1).

**Bompas et al. (2020)** instructed 8 participants to saccade to peripheral visual onsets as fast as they can, and either ignore or stop in response to a central visual onset. The instruction to ignore or stop to the signal was given in separate blocks, inducing measurable pro-active slowing, i.e. an increase in no-signal RT during stop blocks compared to ignore blocks. The signal appeared after the target with SOA 50, 83 and 133 ms, on 40% of trials. The 8 participants were collected in 2 separate experiments, with a total of 8640 and 5472 trials collected per participant. While experiment 2 was a simple blocked instruction design, experiment 1 involved some element of selective stopping: in 5% of trials, a signal of opposite polarity was associated with the opposite instruction (for example: in ignore blocks where most signals were white and to be ignored, a few signals were black and needed stopping). This resulted in a larger difference in mean no-signal RT between the ignore and stop blocks in experiment 2 compared to experiment 1. Data and code from the original article are available [here](https://osf.io/8g9f3/?view_only=83cba9708c5d43fcb25c2e5e6af7d807).

Novel **manual stop-task** dataset. This dataset was collected to quantify the effect of signal brightness on manual non-decision time and to offer a fair test for the decision models (harder task, resulting in slower, less skewed, RT and more errors compared to simple gaze-orienting tasks). It includes two experiments: one ran locally using psychopy on five participants and the other ran online using the pavlovia platform on 45 participants. Participants ran 30 blocks in total. On odd blocks, participants were asked to press the left or right key to quickly indicate whether a centrally presented rectangle was elongated vertically or horizontally. The background was grey, and the rectangle was either light grey (dim condition) or white (bright condition), each contrast being presented for a total of 600 trials. In even blocks, participants were asked to press the left or right key to indicate the side of a peripheral black square, but to withhold their response on detection of a central grey or white rectangle (stop-signal). Dim and bright stop-signals were presented with equal probability on half the trials (600 trials each), with SOAs 17, 33, 50, 67 or 83 ms for the local data, and 50 or 83 for the online data. Participants were asked to perform the task as fast as they could and refrain from slowing down to avoid making errors. Five participants from the online cohort were excluded for poor compliance (low accuracy and infrequent responses showing poor engagement), resulting in 40 participants. All data and code are available [here](https://osf.io/qzywf/?view_only=333db565300c4d75892863bc89bca75d).

### Datasets Analyses

**Mean RT estimates** across all datasets are performed after excluding trials with RT < 0 or > 700 ms and directional errors, if any. The effect of signal properties on mean RT is calculated from specific blocks where those signals were used as targets. The effect of instruction is characterised by contrasting mean baseline (no-signal) RT across the ignore and stop instructions. Datasets featuring only the stop instruction (Boucher et al. 2007a and monkey data) are split into two, according to the nature of the previous trial: trials following signal-present trials (expected to induce more pro-active slowing) versus trials following signal-absent trials. This leads to smaller adjustments than the ignore versus stop comparison, which is why the latter is preferred when available.

**Dip onset time** estimation follows the same procedure as in Bompas et al. (2020), with only minor adjustments to accommodate for differences across datasets. Briefly, RTs for signal-present trials are locked on signal onset and pooled across all SOAs (black line on Figure 1). Histograms are built using bin-sizes of 4 or 3.33 ms for saccadic RT (reflecting the refresh rate of the eye-tracker), and 10 ms for manual RT, then smoothed with a Gaussian kernel to achieve 1 ms resolution. For comparison, a surrogate signal-absent RT distribution is constructed, by submitting signal-absent RTs to the same transformations as the signal-present RTs (i.e. locking these onto hypothetical matched signal onsets, binning and smoothing). The time of maximum interference (T_Max_, red dot on Figure 1) corresponds to the peak of the distraction ratio (N_signal-absent_ – N_signal-present_) / N_signal-absent_, and dip onset time (T_0_, blue dot on Figure 1) is obtained by going backwards in time from T_Max_ until the distraction ratio falls below 2%. Dip time extraction is automatic for all saccadic data with high number of trials. Conditions leading to weaker interference (eg. manual responses under the ignore instruction in Campbell et al. 2017) or relying on a small number of trials (eg. post-signal trials or post-alcohol analyses) are visually inspected and aberrant estimates are removed manually.

**Manual output time** is estimated from the comparison between simple manual and saccadic RT distribution, as described in Figure 2. In addition to a simple fixed delay, three noisy distribution profiles are compared: a uniform distribution (characterised by two parameters, a delay and a range), a Gaussian distribution (characterised by its mean and standard deviation), and a Gamma distribution (characterise by its shape and scale added to a fix delay). The best parameters for each profile are those minimizing the Kolmogorov Smirnov distance between the observed and predicted baseline manual RT distributions. For the fixed delay, the best value is simply picked amongst all the integers from 0 to 300 ms. For the three noisy profiles and each participant, the best parameters are selected following a grid search approach amongst 30,000 simulated noise distributions. An initial round uses large steps between parameter values in order to comfortably cover all participants, and then a finer grid is used to provide better tuning. Because the Gamma distribution has 3 free parameters, while the uniform and Gaussian distributions have only 2, we use larger step sizes throughout when fitting the gamma distribution, to ensure that a similar number of comparisons were performed across the three distributions.

**Group Statistical Analysis.** All reported F and t values relate to repeated measures ANOVA and paired t-test respectively, unless stated otherwise. Null findings are further confirmed using Bayesian statistics, conducted in JASP (JASP Team, 2020). We use equal prior probabilities for each model and 10000 Monte Carlo simulation iterations and report the Bayes Factor for the null (BF_01_), which indicates the ratio of the likelihood of the data under the null hypothesis compared to hypothesis that a difference is present. We interpret BF_01_ of 3 or higher as evidence for the null.

## Empirical Data—Results

In this section, we provide an empirical characterisation of NDT, its sensitivity and immunity to a range of factors, and its suitability for investigating individual differences. Here, empirical NDT refers to dip onset time (T_0_), as obtained from the interference method. **Figure 3** summarises the effect of different factors modulating RT and illustrates which ones do so by modulating empirical NDT (all exogenous factors), and which ones do not (all endogenous factors). **Figure 4** further displays individual data points from each dataset and experiment contributing to Figure 3, and illustrates the individual differences in T_0_, the robustness of which is further illustrated on **Figure 5**. **Figure 6** focuses on manual output time and **Figure 7** provides a summary of all empirical findings.

**Figure 3.**
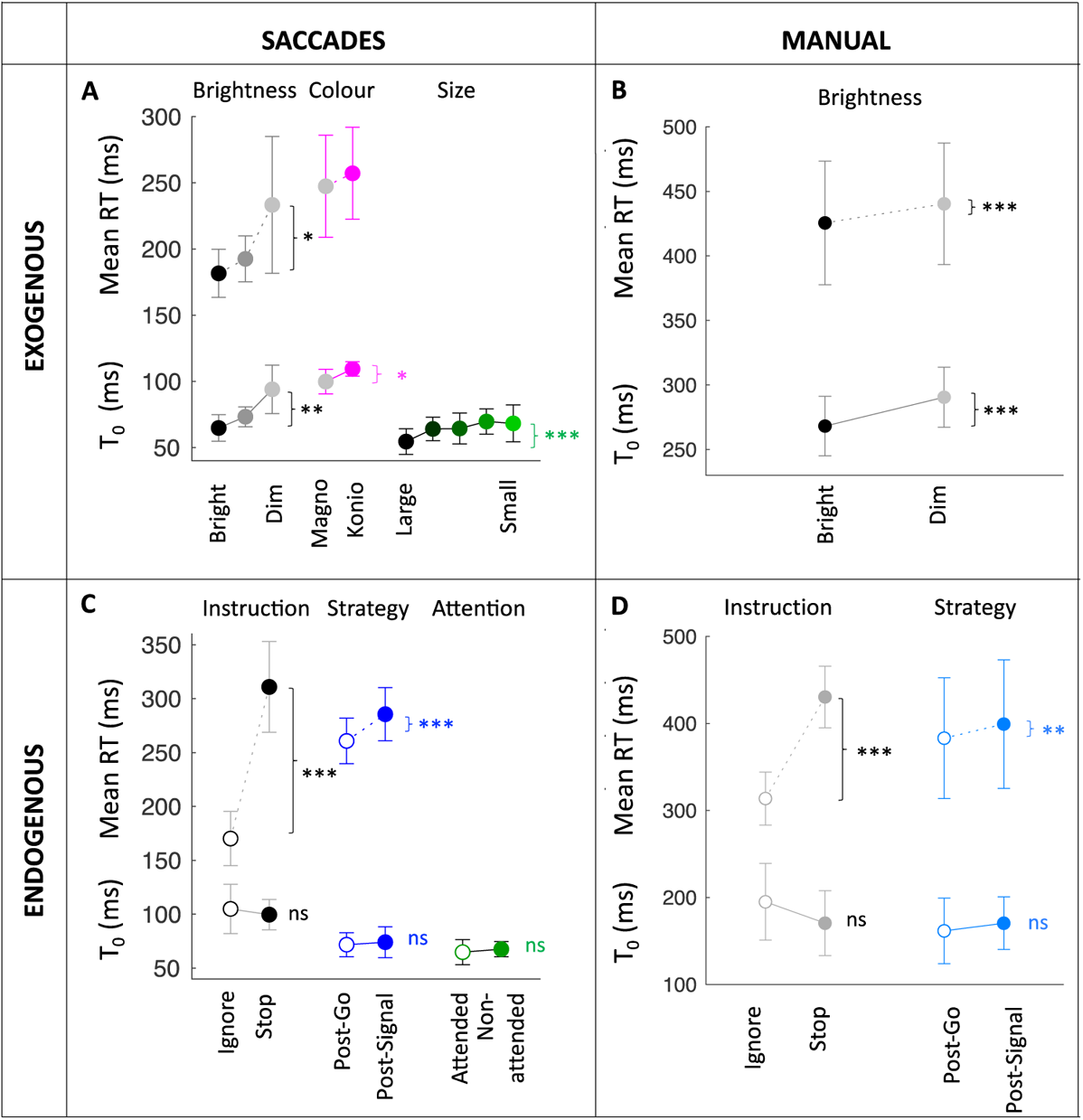
Exogenous factors (top) modulated NDT, endogenous factors (bottom) did not. Data points show group averages, pooled across similar experiments, based on 8 saccadic and 3 manual datasets. Y-axes show average T_0_ (continuous lines), alongside mean RT (dashed lines) when available. Error bars show the standard deviation of the group. Stars indicate statistical significance *p < 0.05, **p < 0.01, ***p < 0.001; ns: non-significant (here all p > 0.2, with BF_01_ > 2). Datasets included: saccades and brightness (Bompas et al. 2009b; free choice dataset); saccades and colour (Bompas et al. 2009a); saccades and size (Buonocore & MacIntosh, 2012); manual and brightness (manual stop task); saccade and instruction (Campbell et al. 2017; Bompas et al. 2020); saccade and strategy (monkey data; Boucher et al. 2007a); saccade and attention (Buonocore & MacIntosh, 2012); manual and instruction (Campbell et al. 2017; Boucher et al. 2007a).

**Figure 4.**
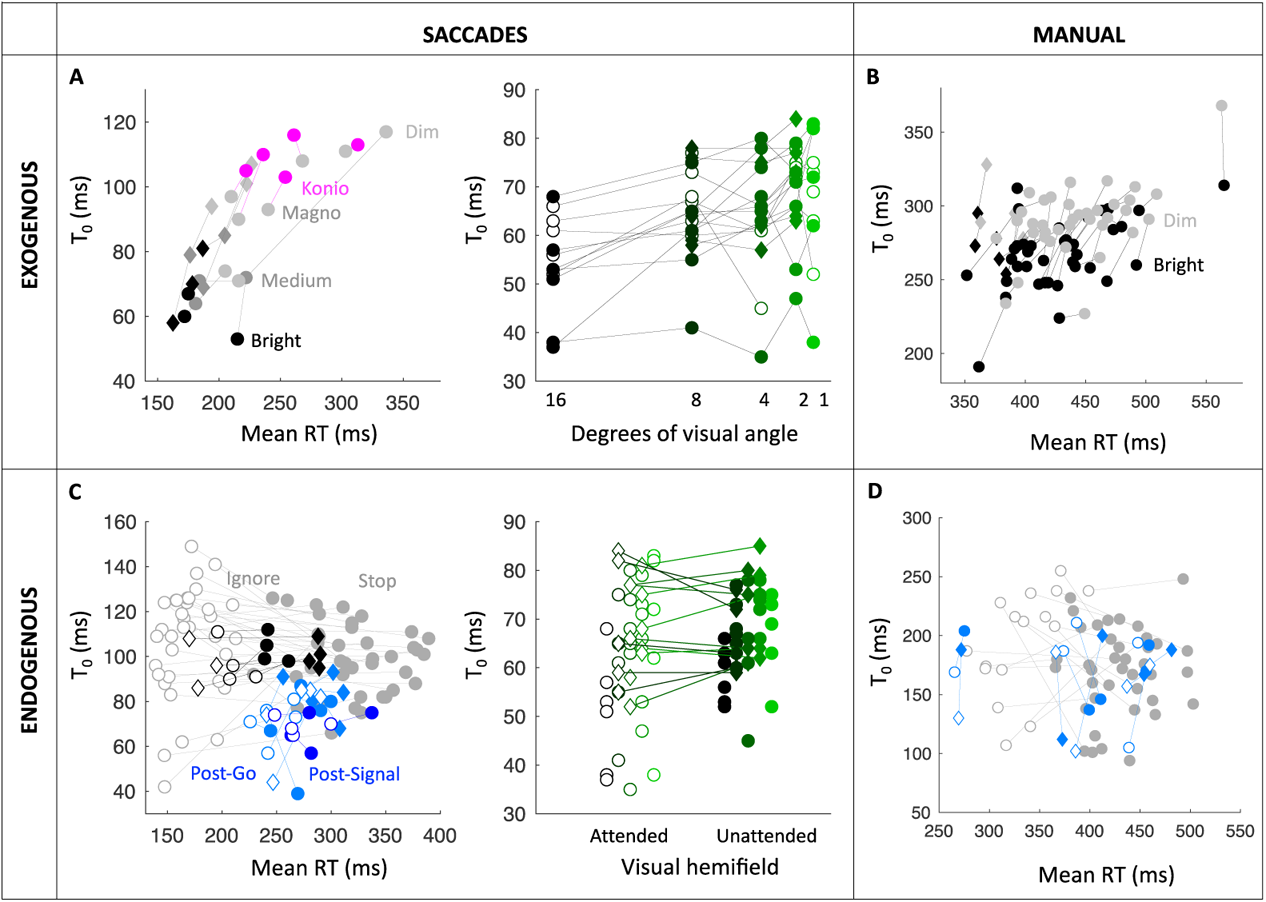
Individual data from each dataset contributing to Figure 3. **A.** Brightness (black to grey circles and diamonds: bright to dim in Bompas et al. 2009b and the free choice data); Magnocellular versus koniocellular (grey to magenta circles: achromatic to chromatic in Bompas et al. 2009a); Size (black to green circles (resp. diamonds): 16 to 1 deg wide stimuli from exp 1 and 2 (resp. exp 3) from Buonocore & MacIntosh, 2012). **B.** Brightness (black to grey diamond and circles: bright to dim from the novel stop-task dataset, local and online experiments). **C.** Instruction (empty to full grey circles: ignore to stop instruction from Campbell et al. 2017; empty to full black circles and diamonds: same from exp 1 and 2 in Bompas et al. 2020); Post-signal slowing (empty to full dark blue: post-signal to post-go trials from monkey data; light blue circles and diamonds: same from unimodal and bimodal conditions in Boucher et al. 2007a); Attention (empty to full green circles: ipsilateral to contralateral distractors from Buonocore & MacIntosh, 2012). **D.** Pro-active slowing (empty to full grey circles: post-signal to post-go trials from Campbell et al. 2017; light blue circles and diamonds: same from unimodal and bimodal conditions from Boucher et al. 2007a).

**Figure 5.**
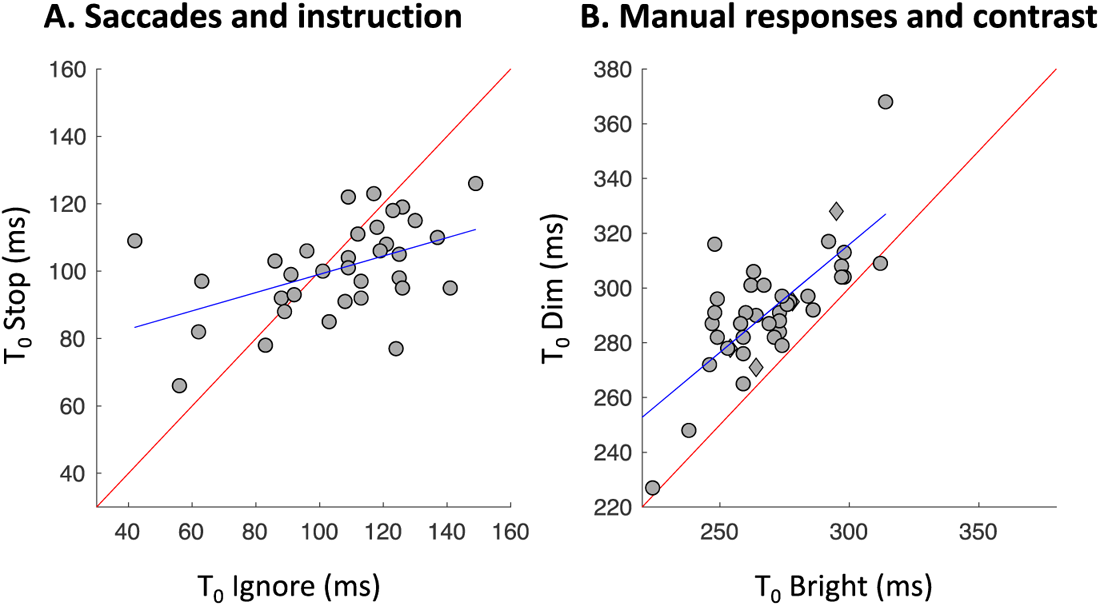
Cross-individual correlations between the T_0_ measured in two conditions. The red line is the identity line. The blue line is the regression line. **A.** Data from Campbell et al. 2017 saccadic blocks for the 34 individuals showing reliable dips in both the stop and ignore instructions. **B.** Data from the novel manual stop-task for the 41 individuals showing reliable dips in the dim and bright conditions, pooling across the online (circles) and local (diamonds) experiments.

**Figure 6.**
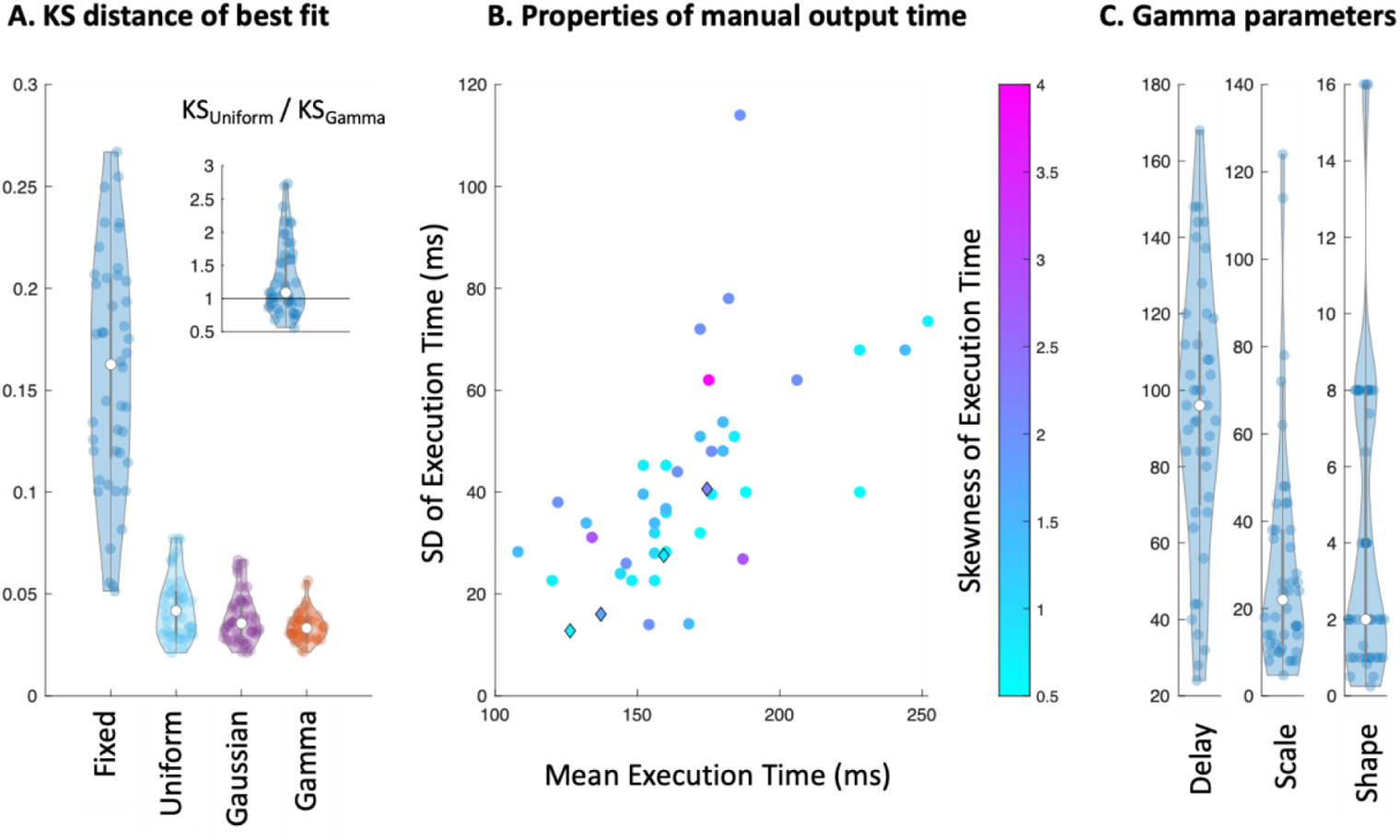
Properties of button press output time distribution, estimated from the distribution that should be convolved with saccadic RT to minimise its distance to the manual RT, as measured with the Kolmogorov Smirnov (KS) statistics. The 44 data points each represent one participant, across two datasets with fast manual and saccadic responses to peripheral targets without distractors (circles: ignore condition in Campbell et al. (2017); diamonds: exp 3 in Bompas et al. (2017)). RT data correspond to fast manual and saccadic responses to peripheral targets without distractors. **A.** KS statistics for the best fitting parameters for each type of motor noise considered, from the worst performing (fixed value) to the best (gamma distribution). The insert directly compares the most commonly used noisy distribution (uniform) to the best performing (gamma). Points show the KS ratio, with values higher than 1 showing better performance for gamma. **B.** Mean, standard deviation and skewness of the best-fitting Gamma distributions across participants. **C.** Best fitting individual values for each of the three parameters of the gamma distribution.

**Figure 7.**
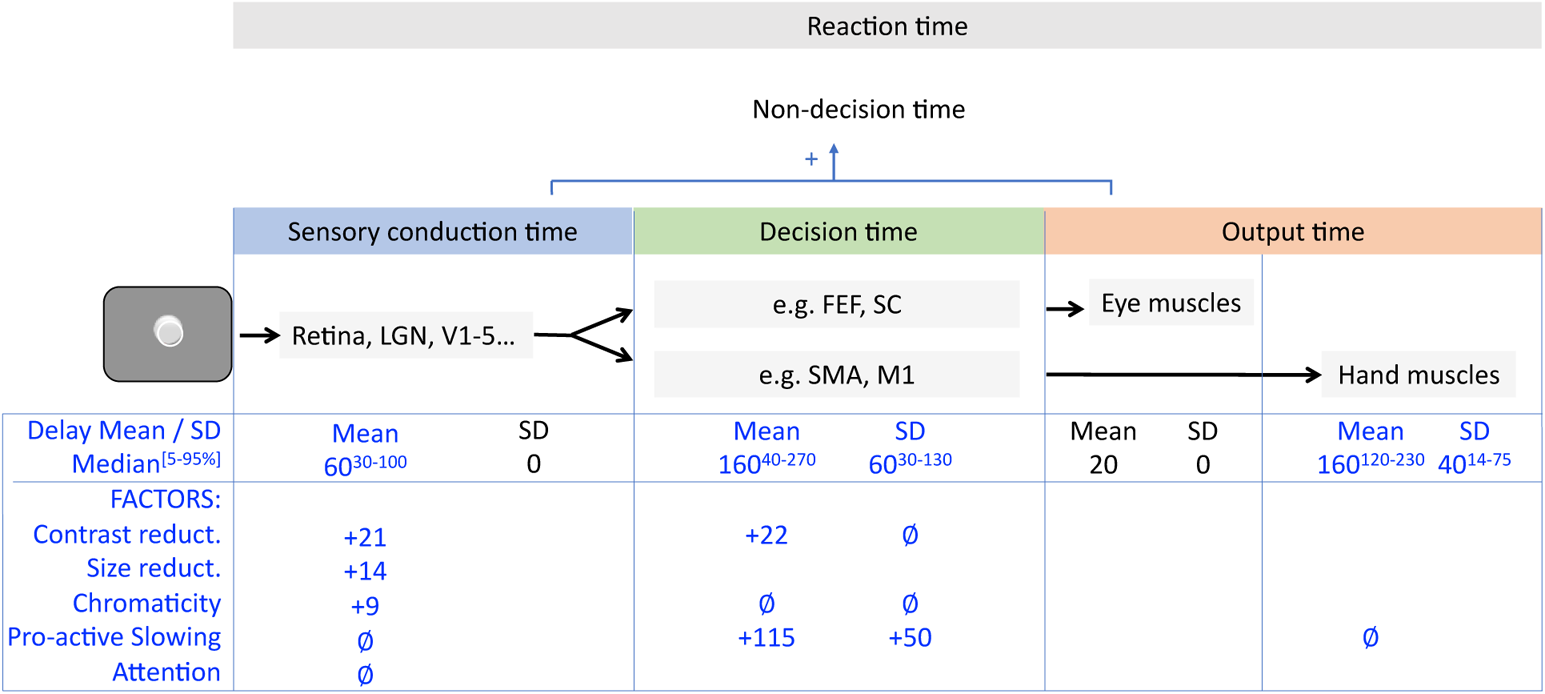
Summary of main assumptions (black font) and conclusions (blue font) from our empirical section. **Top**. Neuronal information flows through visual areas (LGN: Lateral Geniculate Nucleus; V1-5: visual cortex) to action decision areas (e.g. FEF: Frontal Eye Field; SC: Superior Colliculus; SMA: Supplementary Motor Area; M1: primary motor cortex) and to muscles. **Middle.** Mean delays incurred by each stage and across-trials variability. Values indicate the rounded medians across all available estimates (all conditions and participants) and superscript indicating the 5 and 95 percentiles. **Bottom.** Effects of specific factors on the time incurred within each stage and its variability across trials. Here, pro-active slowing refers both to instructions and spontaneous slowing. Values indicate the mean difference across the conditions, when significant (e.g. reducing contrast increases conduction time by 21 ms across relevant datasets, and decision time by a further 22 ms). *Ø* symbols indicate non-significant modulations, with Bayesian evidence favouring an absence of effect. Empty cells indicate aspects that were not tested here.

### Non-Decision Time is Modulated by Exogenous Factors

Low-level visual properties of signals, such as brightness or colour, are obvious ways to manipulate exogenous signals. Neuronal recordings in monkeys performing fast visually-triggered saccades provide evidence that the initial volley of visual activity within areas such as the superior colliculus or the frontal eye field is delayed by up to 50 ms when luminance contrast is highly reduced (Buonocore et al., 2021; Li & Basso, 2008; Marino et al., 2012; Ziv & Bonneh, 2021) and by up to 30 ms when contrast is chromatic rather than achromatic (White et al., 2009). **Figures 3A-B and 4A-B** confirms that empirical NDT measured with the interference method showed this expected pattern, providing an important validation of our approach.

Decreasing brightness resulted in a significant increase in dip onset time. For saccades empirical NDT was on average 65, 73 and 94 ms for high, medium and low brightness (F(2,5)=9.7, p=0.005). For button presses empirical NDT increased from 268 to 290 from high to low brightness, t(44) = 9.6, p<10^−11^). A colour change from signals visible to the magnocellular pathway to signals of equal salience that are invisible to this pathway also led to an increase in dip onset time (on average 100 vs 109 ms, t(4)=3.3, p=0.03), also consistent with previous behavioural literature (Bompas & Sumner, 2008).

Decreasing distractor size led to an increase in dip onset time (on average from 54 to 68 ms for the largest and smallest stimuli, repeated measures ANOVA F(4,36)=6.5, p<0.001). This finding is also expected from the neurophysiology literature, because smaller size has lower total luminosity and also maps onto higher spatial frequency, known to increase the latency of the first visually-driven spikes in superior colliculus (Chen et al., 2018), and to delay saccadic inhibition during reading (Stampe & Reingold, 2002).

Figures 3–4 also display, for comparison and when available, the mean RT observed when these same stimuli were used as targets rather than interferers. Decreasing brightness significantly increased mean saccadic RT (F(1.013,10) =10, p = 0.023 after Greenhouse-Geisser correction) and manual RT (t(44) = 11.9, p<10^−14^), while this effect did not reach significance for colour (t(4) = 2.17, p = 0.1). Similar data was not available for the size comparison.

On saccadic data, because sensory and motor delays are expected to show negligible noise (see Discussion), the visual interference method allows us also to directly assess the effect of visual properties on mean decision time, simply given by mean RT – T_0_. Reducing brightness from medium to low led to a further 20 ms increase in decision time (F(2,10) = 4.3, p = 0.045), in addition to the increase in non-decision time. No change in variability was observed (BF_01_ = 1.35). No change in decision time was observed between equally salient chromatic and achromatic contrasts (t(4) = 0.05, p > 0.9, Bayesian evidence favouring the null with BF_01_ = 2.03).

### Non-Decision Time is Unaffected by Endogenous Factors

Whether spontaneous (following an error or a perceived increase in task difficulty) or instructed, pro-active slowing is one manifestation of the broader construct of speed-accuracy trade-off. Neuronal recordings during saccadic RT tasks show that changes to speed-accuracy settings do not modulate the initial rise of visually-driven neural activity, strongly suggesting these do not affect sensory conduction delay (Heitz & Schall, 2012; Reppert et al., 2018).

In line with this, **Figures 3C and 4C** show that, for saccades, strategic adjustments producing large changes in baseline RT, such as pro-active slowing in response to changes in instruction (t(47) = 18, p<10^−22^) or strategy (post-signal slowing, t(13) = 9, p<10^−6^), induced no changes in saccadic empirical NDT (paired t-test across 4 datasets, on 55 participants including 4 monkeys: t(54) = 0.7, p = 0.47, BF_01_ = 5.9, showing evidence for the null). Particularly striking is that dip onset time was no different whether the signal creating the dip was to be ignored (task irrelevant) or acted upon (very important to the task, p=0.30, BF_01_=3.5), despite an average 154 ms change in mean RT, confirming the conclusions from Bompas et al. (2020), and extending them to trial-to-trial strategic pro-active slowing (p=0.64, BF_01_=3.3).

**Figures 3D and 4D** generalise these conclusions to button presses: instruction had no clear effect on manual dip onset time (t(15)=1.2, p=0.25, BF_01_=2.14), contrasting again with a clear change in strategy evidenced by the mean RT difference between the ignore and stop manual blocks (314 to 430 ms, p<10^−23^). Although the evidence didn’t clearly support the null (BF_01_<3), the pattern of results was similar to that observed on saccadic RT. However, only 16 out of 40 participants showed dips both in the ignore and stop manual blocks. This is mainly down to SOAs in this dataset being too long to be optimal for producing manual dips, especially in the ignore instruction where dips are more transient. This suggests that the stop task could be a better tool for estimating dip onset time in the manual modality.

The Boucher et al. 2007a dataset (light blue series Figure 4D) was also analysed to further investigate the effect of strategy on manual dip onset time. This dataset did not use button presses but joystick orienting responses, either alone (unimodal blocks) or alongside a saccadic response (bimodal blocks), on the same 5 participants. The effect of strategy on dip onset time and baseline RT was analysed in a similar way as for the saccadic modality, by contrasting post-signal and post-go trials. A 2-way RM-ANOVA (unimodal versus bimodal; and post-go versus post-signal) revealed modest but clear spontaneous slowing following signal-present trials (16 ms increase in mean baseline RT, F(1,4) = 29, p = 0.006), but no significant effect on dip onset time (9 ms increase, F(1,4) = 0.2, p = 0.67, BF_01_ = 2.12). These results should be interpreted with caution as trial numbers were low in some conditions (signal present on two trials in a row), which may have further impacted the reliability of T_0_ estimates.

Last, as a different type of endogenous modulation, we also report the lack of effect of visuo-spatial attention on saccadic empirical NDT (green series on panel C), as shown by contrasting the effect of distractors appearing on the attended side of the screen (i.e. ipsilaterally to the target, which always appeared on the right) or the unattended side (contralaterally to the target). This lack of difference was observed first across subjects (contrasting exp 1 and 2 from Buonocore & McIntosh (2012): F(1,8)=0.29, p=0.6, BF_01_ = 77), and then replicated within-subject (exp 3: F(1,6) = 0.005, p = 0.95, BF_01_ = 3.4), both tests showing evidence for the null. Since visuo-spatial attention is often conceived as a boost in salience, this result may appear counterintuitive. However, we keep in mind that this study uses highly salient stimuli, which may therefore be less prone to attention modulations.

### Non-Decision Time Shows Robust Individual Differences

Noticeable in Figure 4 are large individual differences. Do these reflect noise in our estimates, or are these repeatable individual traits? The Campbell et al. (2017) and the novel manual stop-task data both involved one major empirical manipulation and over 40 participants, enough for a first assessment of the robustness of individual differences in dip onset time. **Figure 5** shows that, in both datasets, dip onset times were correlated across conditions. **Figure 5A** shows the correlations between the T_0_ measured from separate blocks under the ignore and stop instructions for the saccadic data (r(34) = 0.48, p = 0.0042; only 16 individuals showed reliable dips in both condition for manual data, so these are not reported). **Figure 5B** shows the correlation between the onset time of dips produced by interleaved dim and bright signals (r(41) = 0.78, p < 10^−9^). One can also note the upward bias on the latter, reflecting the sensitivity of dip onset time to this exogenous factor. No such bias was observed in 5A, again reflecting the immunity of dip onset time to endogenous factors. While the correlation in the saccadic data can only be driven by sensory delay, in the manual case it may be driven both by sensory delay and output time.

### Manual Output Time is Noisy and Often Skewed

In this section, we aim to provide an overview of the main defining features of manual output time distribution and their variations across participants (**Figure 6**). For this, we applied the same logic and methods as in Bompas et al. (2017) on a larger sample of 40 participants from Campbell et al. (2017), focusing on simple reaction times from button presses (see Discussion for generalisation to other response modalities and other tasks).

**Figure 6A** illustrates the success of each noise distribution hypotheses at reducing the distance between the saccadic and manual RT distributions, i.e. at approximating manual output time distribution. Unsurprisingly, the fixed value clearly underperformed compared to the noisy distributions, unable to capture the extra variance in manual RT data. The uniform distribution came second last, losing both to the Gaussian and gamma distributions (t(43) = 3.4, p = 0.001; t(43) = 5.3, p<10^−5^). The gamma distribution did best overall, but only marginally so compared to the Gaussian distribution (t(43) = 1.8, p = 0.077). The insert further shows that the better performance of the gamma distribution compared to uniform was driven by about one third of the participants (KS ratios above 1.5), and this preference was correlated with the skewness of the best-fitting gamma distribution (r = 0.54, p < 10^−3^).

**Figure 6B** illustrates the main properties of the best fitting gamma distribution, including the skewness. Despite the large differences across participants, we note that none of the best fitting distributions across all three hypotheses led to a SD close to zero (as assumed when using a fixed value for non-decision time), nor to a skewness close to zero (as assumed when using a Gaussian or uniform distribution). We therefore conclude that inter-trial variability and skewness are likely features of non-decision time and should constitute desirable parameters for researchers interested in accurately modelling non-decision time. We come back to this conclusion in the Discussion. **Figure 6C** further displays the individual parameters for the best fitting gamma distribution. Exponential distributions (as considered in Ratcliff, 2013) are a special case of gamma distribution with a shape of 1. This corresponded to 9 participants out of 44.

The presence of noise in manual output time means that the relationship between dip onset time and non-decision time is less straightforward for manual compared to saccadic responses. Indeed, manual non-decision time is inherently noisy, but dip onset time is a single value by construction. However, when sufficient trials are available, the expectation is that manual dip onset times should tend towards the minimum value for non-decision time (Bompas et al., 2017). In datasets featuring both manual and saccadic data, this minimum value can be estimated independently for each participant by taking the sum of the sensory conduction delay (saccadic T_0_ – 20 ms) and the minimum value of the manual output time (Delay parameter displayed on Figure 6C). The Campbell et al. 2017 data partly met this expectation: observed manual dip onset times were on average equal to the predicted minimum non-decision time (both were 170 ms), not significantly different from each other (BF_01_ = 5.6). However, the success of this prediction is mitigated by the lack of correlation between observed and predicted values across individuals, which could be due to individual manual dip onset times not being reliable enough for this data.

Our estimates of manual output time pertain to button presses, by far the most popular type of manual response. However, by definition, the main properties of motor output time will depend on response modality. The pathways relied upon by button presses, screen touches, joystick or mouse movement, finger pointing, or hand reaching may overlap (from the motor cortex to arm and finger muscles) but the RT will include different elements of response execution. Researchers interested in decomposing manual output time into a pre-motor and motor execution components may use electromyograms to detect the onset of muscle activation (like in Weindel, 2021). Subtracting this onset time from the RT on each trial reveals the distribution of motor execution time. For instance, performing this analysis on the 6 participants from Gulberti et al. (2014) data suggests that, for finger pointing, the time required for lifting the finger from its starting position is best captured by a Gaussian distribution (mean of 55 ms and SD of 18 ms). Although only 6 participants were available, our analyses suggest those features are highly preserved across the two variants of the task introduced (hand only or bimodal, r > 0.89 for mean and SD) and could therefore constitute stable traits.

### Conclusions from Empirical Data

Figure 7 summarises our assumptions and empirical findings. Note that exact numbers are of little importance here as they will depend on the data used. The key message is that the approach described here allows us to estimate the contribution to RT of each stage of the decision process and its susceptibility (or immunity) to a range of factors.

## Visual Interference Versus Decision Models

### Rationale

Now that we have established that the visual interference method produces estimates that are both reliable and match the expected properties of non-decision time based on neurophysiology, we move on to contrast our approach with the standard approach for extracting NDT from decision models, using the three most popular simple models: the EZ drift diffusion model (Wagenmakers et al., 2007), the DDM (Ratcliff & Rouder, 1998) and the LBA (Brown & Heathcote, 2008).

Our modelling choices aims at addressing three main questions. First, we ask if NDT estimates provided by default (blind) model implementations can be trusted. The general answer being no, we further ask what can be done to improve the models’ outcome. We first adopt an “informed” approach to choosing free parameters, and then introduce a “hybrid” variant, combining the interference method with modelling of the decision parameters. The answers provided here may only apply to the dataset under consideration, but we hope they illustrate important issues and a possible way forward.

We use the dataset that, amongst all the ones presented here, lends itself best to fitting these kinds of model and where our predictions are the most straightforward: the shape discrimination blocks from our manual stop-task dataset, using dim or bright targets. While most previous model validation exercises on empirical data focused on manipulating decision parameters while keeping NDT unaffected (with mitigated success as discussed above, see Dutilh et al. (2019)), ours directly targets NDT while keeping decision unaffected. Our aim is similar to those previous attempts, in that it offers an opportunity to assess the sensitivity, specificity and validity of modelling outcomes. However, in contrast to previous attempts, we have an empirically derived measure of NDT for comparison and stand on firm ground when expecting the other parameters to be unaffected. Indeed, in contrast to a speeded detection task where brightness would affect decision time, the shape discrimination task data that we use here has no reason to be harder on dim than bright shapes, as both are clearly visible and the difference between vertical and horizontal shapes is easy to tell at both levels. Moreover, accuracy and standard deviation of RT are no different on this data – only the mean RT differs, consistent with a specific change in NDT.

To answer each question, we contrasted three variants:

1. A “blind” variant, meant to capture a default approach in the field in the absence of hypotheses (a minimal set of free parameters, all allowed to vary across participants and contrast levels)
2. A “informed” variant, inspired by the spirit of the present article (on this dataset, we expect only NDT to vary with contrast, and trial-to-trial variability in NDT to be present)
3. A “hybrid” variant, which is the same as the blind variant but constrains the model NDT to be equal to the observed T_0_.

### Modelling Methods

Model fits are performed using freely available code and default, non-hierarchical, fitting approaches (all code is available [here](https://osf.io/qzywf/?view_only=333db565300c4d75892863bc89bca75d)). **Table 1** summarises the parameters for each model and variant. We assume readers are familiar with the principles underlying simple decision models and will therefore only provide a very brief overview here. The central assumption common to these models is that decision time is the time taken by a decision variable to progress from a baseline level and reach a threshold. This decision time is added to a non-decision time to form the reaction time. Decision time is determined by the distance between baseline and threshold (boundary separation), and by a drift (or rise) rate which can vary across trials. The LBA possesses only one threshold, towards which all response options (e.g. correct and incorrect) race until one wins. In contrast, the DDM envisages decision as a journey between two boundaries, each coding for alternative response options. Both models assume across-trial (Gaussian) variability in drift rate, but they differ in their assumptions regarding the other sources of variability. The DDM assumes Gaussian within-trial variability in the drift, while LBA assumes a linear rise to threshold. The LBA assumes (uniform) trial-to-trial variability in starting point, in contrast to the standard version of the DDM used here.

**Table 1.** Parameter descriptions and treatment for each model. Free parameters are allowed to vary across participants and contrasts. Some parameters are only allowed to vary across participants but not contrast (Fp). The remaining parameters are scaling parameters or ignored options (grey fonts).

| EZ parameters |  | Blind |  |  |
| --- | --- | --- | --- | --- |
| Non-decision time | $T_0$ | Free | | |
| Drift rate | $v$ | Free | | |
| Boundary separation | $a$ | Free | | |
| Within-trial variability | $s$ | 0.1 | | |
| DDM parameters |  | Blind | Informed | Hybrid |
| Non-decision time | $T_0$ | Free | Free | Given |
| Drift rate | $v$ | Free | Fp | Free |
| Boundary separation / Threshold | $a$ | Free | Fp | Free |
| Across trial variability in $T_0$ | $sT_0$ | 0 | Fp | 0 |
| Across-trial variability in $v$ | $sv$ | 0 | 0 | 0 |
| Within-trial variability | $s$ | 0.1 | 0.1 | 0.1 |
| Response bias | $z$ | 0 | 0 | 0 |
| Across-trial variability in $z$ | $sz$ | 0 | 0 | 0 |
| LBA parameters |  | Blind | Informed | Hybrid |
| Non-decision time | $T_0$ | Free | Free | Given |
| Drift rate toward correct response | $v$ | Free | Fp | Free |
| Across-trial variability in $v$ | $sv$ | Free | Fp | Free |
| Starting point variability | $A$ | Free | Fp | Free |
| Boundary separation | $b$ | Free | Fp | Free |
| Drift rate toward incorrect response | $v_0$ | 1-$v$ | 1-$v$ | 1-$v$ |

Parameters for the EZ diffusion model are calculated using custom code from Wagenmakers et al. (2007). These parameters are also used as initial parameters for the DDM, as part of the default pipeline using the DMAT toolbox in Matlab (Vandekerckhove & Tuerlinckx, 2008). Data are accuracy coded, so that the upper and lower boundaries in the DDM correspond to correct and incorrect responses respectively. Data are fitted by minimising a negative multinomial log-likelihood function, calculated on percentiles of the correct and incorrect RT distribution [10,30,50,70,90,100]). In the blind variant and its hybrid counterpart, we fix all the across-trial variability parameters to 0, following recent expert recommendations to do so to avoid compromising estimates of the main parameters (Boehm, 2018; Lerche & Voss, 2016), and because researchers typically do not have hypotheses about these variability parameters. In the informed variant, we set sT_0_ (the variability in NDT) free across participants, since non-zero trial-to-trial variability in output time is biologically highly plausible in manual data. When this is included, we use (T_0_ – sT_0_/2) for comparison with observed T_0_, reflecting the lower bound of the uniform distribution, in line with our theoretical expectations and analyses of manual dip onset time on the Campbell et al. (2017) data.

The LBA model is fitted via the LBA MATLAB toolbox^3^. Data are accuracy coded, with independent accumulators for correct and incorrect responses. Each participant’s data are fitted independently, from 50 random combinations of starting parameters, and minimising the sum of the negative log likelihood of each observed RT against the model (following the default pipeline in the toolbox; note that we observe excellent parameters convergence across the 50 iterations). As a scaling parameter, we use the standard approach of constraining the sum of correct and erroneous drift rates to 1, and use only one across-trial drift rate variability (sv) for correct and incorrect answers. Because LBA has uniform variability in starting point, we use (b + A/2) in our analyses when referring to threshold. The informed variant of the LBA does not allow across trial variability in NDT, as this option is not yet available for LBA.

### Modelling Results

Throughout this section, we define success by the ability for NDT model estimates to mimic observed T_0_ values, i.e. show:

A. Accuracy (stay close to observed T_0_ on average)
B. Validity (correlate with observed T_0_ across individuals)
C. Sensitivity (decrease with contrast)

A fourth success criteria is specificity, and relates to decision parameters (while the other three relate to NDT): we expect drift rate and threshold to be immune to contrast (see rationale section above), and not show extreme correlations with NDT or each other. Although some correlations across parameters may be genuine within the population, large correlations are more likely diagnostic of trade-offs, and the increases in variance these trade-offs induce in individual estimate reduce their interpretability.

### Default Model NDT Estimates Do Not Correspond to Empirical NDT

Each model was first fitted blind to the manipulation at play, which tends to be the default approach in the field (acknowledging that some studies have constrained models depending on task and manipulation). **Figure 8** summarises the results of this endeavour showing the T_0_ estimates from the best fit for each model variant, against the observed T_0_ from the interference method. The blind columns of **Table 2** further assess model’s performance across all four criteria.

**Figure 8.**
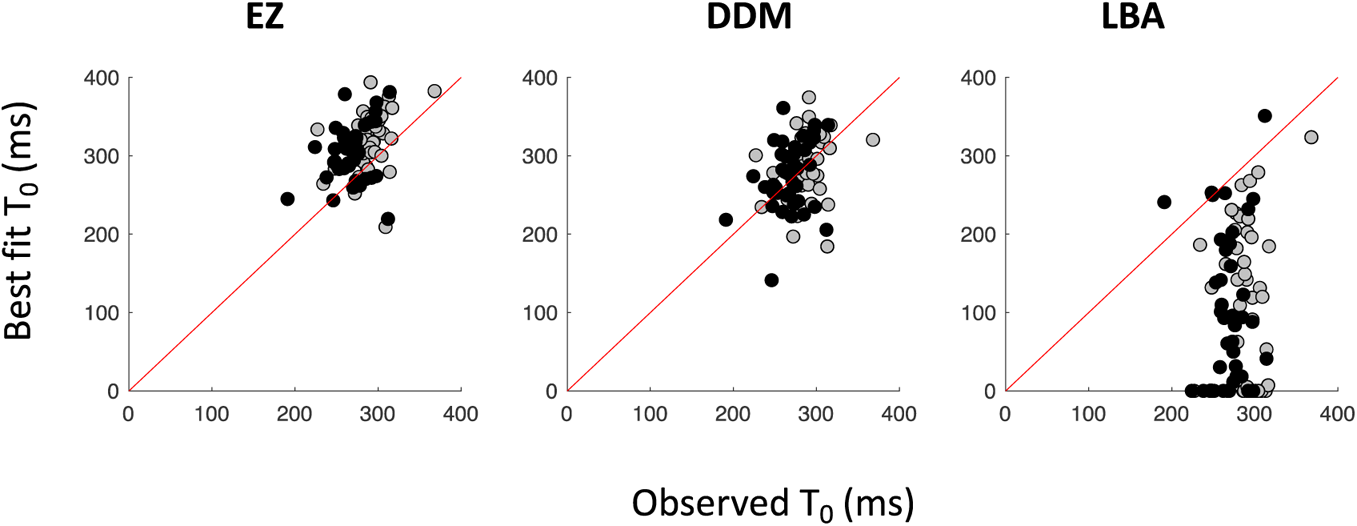
NDT model against empirical estimates for the blind implementation of three common simple decision models. Data points show individual estimates from the novel manual stop task data, using dim (grey) and bright (black) stimuli. The red line is the unity line. Clouds of points centred on the red line (like DDM) indicate good accuracy, while those centred above (EZ) or well below (LBA) indicate systematic biases. Clouds of the same colour oriented diagonally would indicate validity (correlation between observed and model estimates), but none did clearly so. Grey clouds centred right of black clouds indicate contrast sensitivity (EZ and, to a lesser extent, DDM).

**Table 2.**
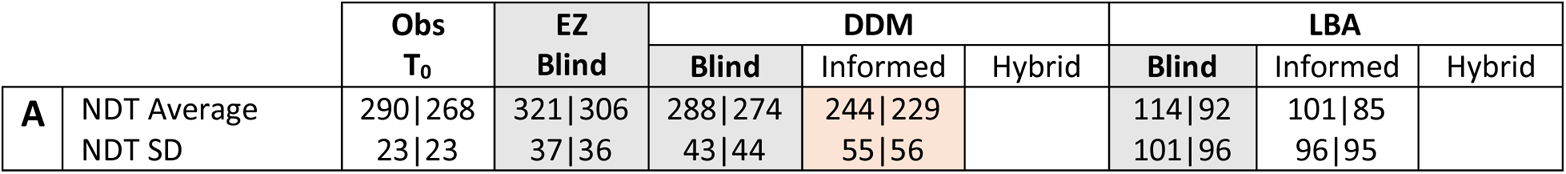
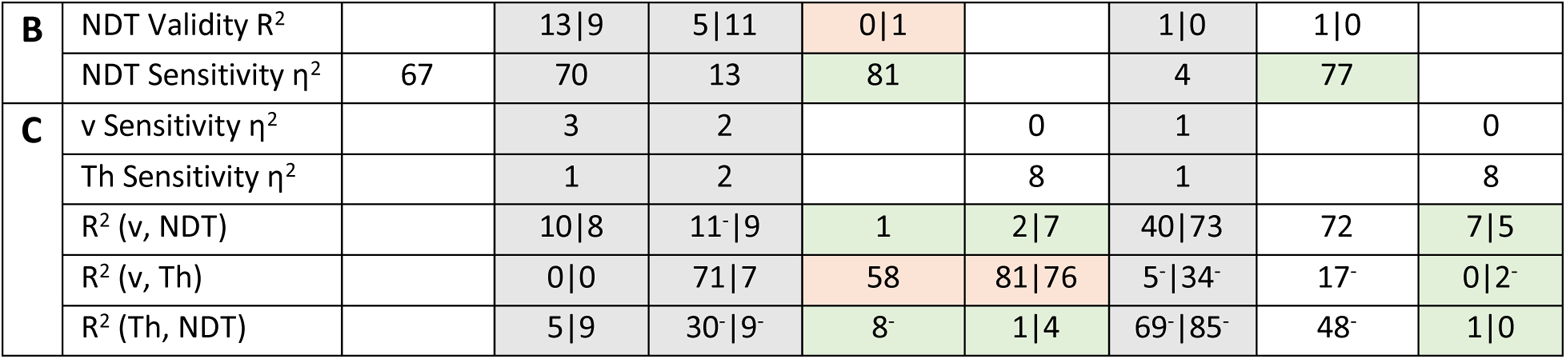
Summary of model performance, alongside observed T_0_. When applicable, values are provided at each contrast level (dim|bright). **A.** Group mean and standard deviation of NDT in ms, reflecting the accuracy criteria (all significantly different from observed except the blind DDM, all p<0.001). **B.** Validity: R^2^ of the correlation between observed and model estimates. Sensitivity: η^2^ of the effect of contrast. **C.** Sensitivity of drift rate (v) and threshold (Th), and correlations across parameters (here high values now indicate poor performance, addressing the specificity criteria). (^−^) indicate negative correlations. **A-C.** Grey shading indicates the blind variants; green and orange shading highlights areas of improvement or deterioration from the blind to the informed or hybrid variants, when the significance of the effect cross a boundary (non-significant, significant at 0.05, significant at 0.001), which on this sample corresponds to effect sizes below 10, between 10 and 30, or above 30.

#### Accuracy

Only DDM was accurate, EZ produced inflated though still reasonable estimates, while LBA showed extreme undershoot. The variability of NDT estimates across participants were all inflated, with EZ closest to observed SD values, followed by DDM and LBA. For LBA, 53% of NDT estimates fell below 100 ms. Considering the first signs of visual activity in the brain occur at the very least some 20 ms after the visual event, and that manual output time in our analyses never fell below 100 ms, any estimates under 100 ms are likely impossible neurophysiologically.

#### Validity

EZ and DDM estimates showed at best modest correlations with observed T_0_ (shared variance of 10 and 8% on average), while LBA showed none. From the above, models in their default implementation appeared unable to accurately capture individual differences in NDT.

#### Sensitivity

EZ showed excellent sensitivity to contrast (comparable to observed T_0_), while the DDM showed only modest sensitivity (just above significance at p=0.05) and the LBA none.

Overall, the simplest model of all, EZ, did comparatively best (despite its poor validity). Its overshoot (by on average 35 ms) could even be argued to be a sensible expectation, for those who consider that the decision in this task is about pressing the left or right button, while determining if the shape is vertical or horizontal belongs to perceptual processes and therefore NDT (we come back to this matter of definition in Discussion). Since T_0_ would only reflect sensory delay and motor output time but not perceptual processes, they would be lower than model estimates. Its specificity was also excellent, with decision parameters showing no sensitivity to contrast and no correlations across parameters. This makes EZ a great choice for researchers who are not interested in individual differences and just want to test hypotheses about experimental manipulation of NDT. In contrast, DDM and LBA both displayed concerning levels of cross-parameter correlations, which could contribute to the large inter-individual variance in NDT estimates. Furthermore, the immunity of their decision parameters to contrast cannot indicate specificity, given that their NDT parameters also showed poor sensitivity to contrast.

### Informed constraint is not enough

Next, we ask whether using a-priori knowledge to guide modelling can improve performance. Our “informed” variant (see Methods for details) only allowed NDT to vary with contrast (for DDM and LBA) and further allowed variability in NDT (for DDM). As shown in Table 2, contrast sensitivity was greatly increased for NDT in both models (η^2^ > 77%) but, crucially, correlation with observed T_0_ across individuals (validity) wasn’t. Although the improved sensitivity was reassuring, it is highly expected (since only NDT was able to vary with contrast), and mainly uninformative unless accompanied by increased validity (since it already formed part of our hypotheses). Accuracy remained very poor for LBA, and decreased for DDM. For the DDM, the SD of NDT values across participants also increased further away from observed values. This may be partly explained by the fact that NDT estimates now result from a combination of two parameters (T_0_ and sT_0_), since the SD of T_0_ - sT_0_/2 is mathematically higher than the SD of T_0_ or sT_0_.

Shifting focus to decision parameters, some of the cross-parameters correlations were reduced, but not systematically so. Because the decision parameters did not vary with contrast, we cannot assess their sensitivity to contrast. In conclusion, our attempts at better constraining parameters, while remaining within the realm of default practices within the field, proved largely unsuccessful. Future research could determine the generalisability of this conclusion to other datasets and explore the potential benefits of using noisy NDT profiles, including non-uniform and skewed distributions.

### Combining Modelling with Empirically Derived NDT Values Helps

Last, we bypassed the attempts to extract NDT from model fits, and instead inserted the individual observed T_0_ values at each contrast level when fitting the models. This “hybrid” variant was therefore, by construction, only informative about decision parameters. Table 2 shows an overall benefit to the LBA, reducing correlations across all parameters (all R^2^<0.07, all non-significant). These improvements may be driven by the forced reduction in decision time (because LBA previously underestimated NDT), leading to an increase in drift rate and a reduction in threshold and its cross-individual variability. As for DDM, the correlation between decision parameters and NDT were reduced, but the correlation between drift rate and threshold remained extremely high (> 70% shared variance). In conclusion, the hybrid method appears to have cured the LBA of the trade-offs that plagued its decision parameters, but its efficiency was only partial for the DDM. Of course, the real test for this approach is to assess whether the hybrid method can result in higher external validity for decision parameters. This can be the purpose of future research.

## General Discussion

### Results Overview

Using an empirical method for characterising sensory delay and motor time in visuomotor decisions, we confirmed the influence of visual properties (luminance contrast, colour and size) on sensory delay and found evidence against an influence of top-down factors such as pro-active slowing and attention. Then, contrasting simple saccadic and manual RT, we delineated the main properties of manual output time, showing that using a fixed value to capture non-decision time provides poor fit to data overall. Furthermore, one third of participants further benefit from skewed distributions over the more standard uniform distribution.

In the last section of the article, we contrasted the conclusions reached from our empirical method with the standard approach using decision models. The EZ model was best suited for detecting experimental manipulations of NDT (while the most popular DDM did poorly and the LBA very poorly). Our results also suggest that models cannot be relied on if researchers are interested in capturing individual differences in NDT, irrespective of modelling choices. We then introduce a novel hybrid approach, which combines the interference method to estimate NDT, with standard modelling to estimate decision parameters. This approach shows promising improvements, through reduced trade-offs across parameters.

### Where Sensory Delay Stops and Decision Begins

Non-decision time is essentially a decision-centric concept, created by researchers interested in modelling decision, to account for delays occurring before and after the decision process at the heart of the model. So, to define non-decision time, we need a conceptual definition of when a decision process starts and ends. Throughout this article, with our focus on speeded visuo-motor tasks, we have defined decision as a process that starts when the first volley of visually triggered activity reaches one of the areas involved in the decision, and stops when a point of no-return has been reached within some neural population coding for an action plan. This definition should be consensual when the task simply involves detecting or orienting to a single target, because the first volley of visual activity is indeed task relevant. But are all scenarios similarly straightforward? For tasks where a perceptual process is required (e.g. is the shape vertical or horizontal) prior to selecting the appropriate action plan (press the left or right button), one may ask whether the delay incurred by this perceptual process should be part of NDT or decision. Below, we argue for the latter, allowing the definition of NDT to remain the same for all speeded visual tasks.

Monkey neurophysiology shows that sensory information flows through to brain areas associated with action decisions, and to areas involved in perceptual processing, all in parallel. In monkey FEF, for example, the first volleys of activity only reflect stimulus onset or offset, while later activity increasingly differentiates task-relevant sensory features, such as colour, shape or motion (see for instance Purcell et al., 2010 for a review). More sophisticated perceptual processing may lead to longer lag before neural activity starts to reflect task-relevant features, but information would still flow directly through as it is processed, rather than wait for a perceptual stage to complete before carrying on to neurons involved with action decision. In computational terms, we see decision as the dynamic integration of multiple signal sources, that become increasingly informed by the task as the perceptual processing evolves. Categorising the perceptual process as part of the decision process also allows us to conceptualise perception generally as making decisions about sensory inputs.

Pragmatically, this proposal also sets a clear distinction between empirical factors that are objective (also referred to as exogenous, automatic or task-unrelated, such as stimulus properties) versus subjective (i.e. endogenous, such as task instructions, or spontaneous adjustments within observers). This distinction is powerful because it provides simple guidelines that the former category affects sensory delay while the latter does not, and these can be used to generate clear predictions when generalising from known empirical results to new combinations of empirical factors. In contrast, if we were to incorporate task-relevant perceptual processing into the definition of sensory delay, both non-decision time and decision time could differ for every empirical design. Indeed, even when a decision network of interest can be defined, such as the saccade generating network (which includes the FEF, SC, parietal cortex and basal ganglia), there are multiple early sensory pathways that feed this network, each with different properties and temporal characteristics, and multiple routes from these early pathways through different visual processing areas that will be more or less relevant for each task (see for instance Bompas & Sumner, 2008). Instead, our proposal is that sensory delay should only vary with exogenous properties but will remain the same whether the task is, for example, to press a button as soon as a string of symbols appears on a screen, or make a discrimination based on their visual, orthographic or semantic features. In turn, this implies that decision may include multiple processes.

### What do Decision Model Parameters Really Mean?

In contrast to the complexity of neural pathways and networks supporting even the simplest decisions (such as making saccades), decision models must be highly simplified to be constrainable and, ideally, theoretically interpretable. One attractive possibility is that the NDT parameter of decision models incorporates perceptual processing as well as the basic sensory delay, and this could explain why reported NDT values are varied and often influenced by strategic top-down factors (see our introduction). This explanation might apply where model-fitted NDT values are much longer than in our empirical technique and monkey cell recordings. But it would not apply where models produce too-short NDT values, and nor would it explain why model-fit NDT can vary so much between highly similar tasks or different experiments with the same task. Therefore, we believe a more productive way forward is to take a brief journey back into the conceptual mathematics of fitting models to response distributions, stripping away the parameter labels we have grown accustomed to.

Three key aspects of human decisions inspired the use of accumulation models: even for the simplest tasks, responses show variance, skew and errors (Carpenter & Williams, 1995; Van Zandt & Ratcliff, 1995). Two of the key parameters of an accumulation-based model (gain and threshold, or drift rate and boundary) provide these outcomes in distributions that approximate the shape of behavioural distributions. These parameters engender a systematic relationship between variance, skew and errors, but behavioural response distributions often do not exactly fit this relationship. Therefore, a parameter (or more) is needed to soak up delay and/or variance without affecting (or with a different relationship with) skew and errors. NDT does (part of) this job in decision models. When it is fixed, it can only account for a simple offset of the distribution, but when it has uniform noise, its value is an indication of the degree to which the simulated response distribution needs non-skewed variance to fit the behavioural distribution (the value will also be influenced by what other parameters exist to help with this, and with the relationship between correct and incorrect responses). We propose to stop assuming that the parameter traditionally called NDT in models maps directly onto sensory and motor delay, and therefore deserves to be interpreted functionally as non-decision time.

If we step back from associating model-fit NDT only with sensory and motor delays, and boundary with caution, one may wonder how simple accumulation models may still help understand cognitive functions. It is worth noting that models provide a valuable service that is often overlooked: they effectively provide smoothing and outlier rejection in a systematic way based on the expected shapes of response distributions (even if you ignore all theoretical associations with their parameters). However, to allow models to help beyond this rather pragmatic purpose, one will need to think in terms of what processes should produce more or less skew, and what processes would inherently affect error rates and which might not. For example, if strategic factors change the NDT parameter, this indicates a change that affects the distribution skew or error distribution in a way that doesn’t quite fit with what boundary or drift rate changes do, at least not under their current implementations in simple models. There are several reasons why skew might not fit the pattern produced by the accumulation of a single variable affected by Gaussian noise, for example if multiple streams of input with distinct strengths or temporal profiles contribute to the decision accumulation or if there is significant leakage (like in the DINASAUR model, see Bompas & Sumner, 2011). We suggest that future research will only address this question if we resist assuming we already know the answer–that changes to NDT always indicate changes to sensory or motor delay.

### No Effect of Strategic Adjustments

A major, apparently generalisable, principle across our results was that NDT is changed by stimulus characteristics but not by strategic modulations, such as proactive slowing associated with higher error risk, real or perceived. This clean difference aligns with monkey electrophysiology literature using saccades but stands in contrast to results from decision models on manual data, as elaborated below.

Our results converge with single cell recordings in the FEF of monkeys performing saccadic RT tasks while being rewarded primarily for speed or accuracy (Heitz & Schall, 2012; Reppert et al., 2018). Visual latency was not affected by reward structure, while the baseline firing rate of peripheral FEF neurons and the gain of the visual response were affected, leading to longer decision time under accuracy emphasis. Note also that the change in gain is at odds with the original theoretical expectation of simple decision models, where threshold is normally conceived to capture caution (an assumption not necessarily supported by empirical results, as described by Hedge et al., 2020). Indeed, combining the above neuronal findings with a simple decision model, Servant (2019) showed that, while a change in threshold alone was sufficient to capture the behavioural performance of monkeys under speed versus accuracy conditions, it could not account for the response of visually-responsive FEF neurons (see also Cassey et al., 2014 for modelling implications of these findings).

For manual responses, our results also indicate that non-decision time is immune to pro-active slowing. This conclusion contrasts with previous observations using simple decision models, in particular the DDM, where changes to the NDT parameter often accompany the expected changes in threshold (Dutilh et al., 2019; Lerche & Voss, 2018). This led some authors to conclude that the standard speed-accuracy manipulation in human lacks specificity, but could also indicate that the models are unable to specifically capture the mechanisms at play (Evans, 2021). Interestingly, some have argued that the effect of speed-accuracy manipulation on manual non-decision time is actually expected (Servant et al., 2021), if one assumes decision activity carries on growing after the decision threshold is reached, with motor execution under the control of a second (later) threshold. However, given our results, we believe the main reason for the discrepancy between our findings and those obtained from modelling is that, in contrast to our empirical approach, decision model NDT parameters are not solely extracting the original conception of what NDT is: sensory and motor time.

### External Validations of Decision Models

Our modelling conclusions may also appear at odds with previous modelling work, in particular parameter recovery articles on simulated data. First, although EZ has been previously shown to be superior to the full DDM for detecting simulated experimental manipulations, this benefit was limited to decision parameters while NDT differences were detected equally well across the two models (van Ravenzwaaij et al., 2017). Note however that this previous work relates to between-group differences, while our results concern within-participant manipulation. Second, our conclusion that models are poor at capturing individual differences in NDT contrasts with previous studies showing that NDT can be recovered quite well in simulated data (Lerche & Voss, 2017). This divergence illustrates the need to go beyond parameter recovery and address the external validity of models to break the deadlocks that currently limit their use and interpretation (Evans & Wagenmakers, 2020). A common goal of cognitive modelling is to *quantify* the cognitive processes underlying behaviour. Although it has been shown that different models can lead to similar conclusions when testing experimental manipulations in behavioural data (Donkin et al., 2011), it is often the case that different models can lead us to very different estimates of these quantities. In our data, estimates of NDT from the DDM were 150-200ms longer than estimates from the LBA, which in isolation could naturally lead us to different assumptions about what the brain is doing during simple decisions. There are limits to our ability to identify superior models using traditional approaches to validation. Different decision models can fit empirical data and capture manipulations equally well, despite different assumptions and architectures (Donkin et al., 2011; Teodorescu & Usher, 2013; Van Zandt & Ratcliff, 1995). External validation of model parameters gives us another benchmark by which to compare our models.

It is worth stressing that parameter recovery is a very different way to judge a model than biological plausibility. Simple forms of the models have been shown to be quite successful by the criterion of parameter recovery (e.g. Donkin et al., 2011; van Ravenzwaaij & Oberauer, 2009). Models that incorporate additional parameters, such as trial-to-trial variability, leakage or conflict parameters in the evidence accumulation process are a step closer to likely biology, but show poorer recovery without additional constraints (Boehm, 2018; Hedge et al., 2022; Lerche & Voss, 2016; Miletić et al., 2017; White et al., 2018). A case in point here would be the incorporation of skewed motor output noise. We concluded above that manual output noise is likely skewed, but incorporating this into a model would risk poorer parameter recovery, given that there would now be two sources of skew (the evidence accumulation and the motor output). The only way to make models tractable is to incorporate simplifications or constraints (Miletić et al., 2017), guided by a clear definition of what the modelling exercise is aiming to achieve. In contrast, if the aim is to include a biologically plausible representation of motor output time, it should probably be skewed. A fruitful way to incorporate constraints is to measure some parameters externally to the model fitting process. Our hybrid modelling here demonstrates that this approach can potentially aid the interpretability of parameter estimates.

### Recommendations For Estimating Sensory Delay

The estimation of sensory delay from the interference approach is in principle amenable to any datasets whereby some stimuli are occasionally presented close in time to a saccade-triggering stimulus (Reingold & Stampe, 2002) or the likely initiation of an endogenously driven saccade (e.g. in reading: Reingold & Stampe, 2004; anticipatory saccades: Salinas & Stanford, 2018; or free viewing: Stampe & Reingold, 2002). Most studies considered here present a peripheral visual target followed, on some trials, by another visual stimulus, but other designs may be equally suitable. For instance, in our free choice task, the chosen stimulus plays the role of a target, while the unchosen stimulus acts as an interferer. The presence of a peripheral target is also not mandatory and can be replaced by any stimulus onset that triggers a saccade (e.g., the onset of a central arrow pointing to a peripheral placeholder, an anti-saccade target, or an auditory cue prompting a saccade to a memorised location), or indeed completely omitted like in free-viewing or reading. Likewise, the interferer doesn’t need to be visual, although the visual modality is the most effective for disturbing saccade plans. For the extraction of dip onset time to be reliable, it is desirable to use enough trials to obtain smooth baseline RT distributions. Although this has not been systematically assessed, in our experience, 100 for baseline and 100 for signal-present is likely a bare minimum. It is also critical to choose SOAs wisely, to avoid producing dips at the edge of the baseline distribution, as these can be more easily confounded with noise. In speeded conditions when the target and interferer have similar salience, the optimal SOA will likely be around 40 ms. However, distractors that are more salient than targets will require longer SOAs, and vice versa. Also, when participants slow down (like in the stop task), SOAs need to increase proportionally. Adjusting SOA as a function of participant mean RT can prove effective, as is common in the stop-task and saccadic inhibition literature.

### Recommendations For Estimating Manual Output Time

The estimation of manual output time illustrated here requires the direct comparison of manual and saccadic data on the same participants and task. Not any task is suitable though. The analyses presented here are conducted on simple RT data in an easy task. In contrast, stop-task data are not usable for our purpose, as they produce baseline saccadic RT distributions that are systematically *more* variable than the manual distributions. This is in stark contrast with simple RT data, where the opposite pattern is robustly found. This suggests that participants adopt different strategies when responding with the eyes or the hand in the context of the stop-task, which violates the premise of our empirical approach. Previous work suggested that one strategy developed by monkeys and humans alike to avoid frequent errors in the saccadic stop-task is to hold on fixation stronger to delay saccades (Lo et al., 2009), a strategy not available to manual responses. This strategy may take resources and be deployed mainly when the perceived risk of error increases, leading to the increased variability compared to the ignore condition where this strategy is never necessary. Similarly, any factor that may affect one modality more than another (e.g. fixation gap effect) is discouraged. Researchers interested in implementing our approach are encouraged to acquire blocks of simple saccadic and manual responses, either together (one saccade and one button press on each trial), or in a blocked design. Based on our experience, randomly alternating saccadic only and manual only trials should be discouraged, as the risk of producing saccades on manual only trials is high.

Regarding interpretation of our findings, our approach relies on the assumption that the added variance in the manual compared to the saccadic modality all originates at the motor stage, rather than at the sensory or decision stages. Although visual pathways are partly shared across modalities, we know that the weighting of the different pathways (magnocellular, parvocellular and koniocellular) into action and perceptual decisions differs across modalities (Bompas & Sumner, 2008). Moreover, these visual pathways project onto different decision brain areas (as depicted in Figure 7), providing ample reasons for the decision process to differ. Despite all this, we previously concluded that saccadic and manual decisions are indeed similar enough to support this approach (Bompas et al., 2017). In this previous work, we compared observed data with predictions from alternative models across a range of measures, including the strength and timing of the interference effect at various SOAs. This work showed that allocating the added variance at the motor stage provided the best match to our data. However, this work was conducted on four participants only, and could therefore benefit from a validation on a larger sample.

### No Trial-to-Trial Variability in Saccadic Non-Decision Time?

Equating saccadic dip onset time with non-decision time relies on variability in non-decision time being negligible, which requires both sensory conduction delay and saccadic output time to be highly consistent across trials. If this assumption was wrong, dip onset time would tend towards the lower bound of non-decision time (providing many trials are available), as it does in the manual modality, with only minor consequences for our conclusions. We address this assumption below.

#### No variability in saccadic output time?

All studies relying on neuronal recordings in FEF or SC during saccade tasks consider output time to be within 10 to 20 ms, and average firing rates over this time window have been calculated to estimate the decision threshold (e.g. Heitz & Schall, 2012). However, this range doesn’t imply that output time is thought to be variable across trials within the 10 to 20 ms range. Rather, it is our understanding that the range may reflect 1) the need to define a time window for averaging neuronal responses and 2) methodological differences across studies. For instance, microstimulation studies in the SC designed to *perturb* saccade trajectory often provides estimates around 10 ms, while 20 ms would be required when the aim is to *initiate* a saccade when premotor burst neurons are inhibited by omnipause neurons, as is the case when triggering a saccade while fixating (Miyashita & Hikosaka, 1996; Munoz & Wurtz, 1993). In our work, we use 20 ms, consistent with the latter scenario and with our earlier modelling work, inspired by recordings in visuomotor neurons in the SC (Trappenberg et al., 2001), and still in line with recent assumptions in the field (Buonocore et al., 2021).

#### No variability in sensory conduction delay?

We know this assumption to be an approximation. Our confidence that this approximation is reasonable in the context of our data derives from indirect evidence, reading across a range of single-cell recording studies measuring neuronal response to visual stimuli in monkey FEF and SC. Based on these, it is our understanding that the approximation holds for salient, low-spatial frequency stimuli most common in this field (0.5 to 2 cycle per degree Gabor patches, bright squares or circles of about 1 degree of visual angle). In particular, some articles would directly contrast the evolution of firing rates in visuo-movement neurons averaged across slow versus fast RT trials. They report the same timing for the initial visually-driven response (e.g. Fig 1C in Ray et al., 2009, showing visuo-movement neurons in FEF). This shows that, if variability exists, it is not sufficient to drive clear differences in reaction time across trials. Other articles would display, for one example neuron, spike time series across many trials (e.g. Figure 2 in Buonocore et al., 2021; or Figure 1 in Chen et al., 2018, both based on SC recordings), again suggesting high consistency of visually-driven neuronal response across trials. However, some variability may be expected when using low-salience stimuli. For instance, Lee et al. (2010) showed that the latency of the first visually-evoked spike in V1 is predictive of saccadic reaction time across trials. This implies that this latency *can* be variable across trials under some conditions. In our data, low contrast stimuli lead to higher RT variability, but this trend was not significant. This pattern is expected though, since an increase in the variability of neuronal latency is reported as early as the retinal level (Bolz et al., 1982). Note that the literature using noisy stimuli (e.g. random-dot kinetograms, Gold & Shadlen, 2000) is not directly relevant to our question here, as the variability is built into the design rather than reflecting properties of the visual pathways.

### Biological Underpinning of Individual differences in Non-Decision Time

We have argued that empirically measured NDT reflects the first volley of signals into the decision process, plus the motor time beyond a decisional point-of-no-return (the point after which the action is going to happen, even if it can be modified in flight). It might therefore be considered surprising that such basic transmission times show marked individual variation. One possible place to start looking for biological underpinnings of such differences is in the white matter tracts. Several studies have reported that speed positively correlates with measures of white matter “integrity” (e.g. fractional anisotropy, FA) in multiple tracts (Haasz et al., 2013; Kuznetsova et al., 2016; Penke et al., 2012; Penke et al., 2010). Correlations between white matter and trial-to-trial variability in RT have also been observed, both in specific tracts and at a whole brain level (Booth et al., 2019; Fjell et al., 2011; Kievit et al., 2016; Tamnes et al., 2012). Further studies have used simple decision models to examine associations with microstructure properties (Forstmann et al., 2011; Imms et al., 2021; Karahan et al., 2019; Yang et al., 2015). Two of these report associations with the NDT. Yang et al. (2015) used a letter discrimination task and reported a correlation between the DDM NDT parameter and radial diffusivity in the body of the corpus collosum. Karahan et al. (2019) used a simple manual RT task and focused on the microstructure properties of task-relevant tracts only, i.e. the optic radiation and the cortico-spinal tract. They reported no correlation with mean RT, but some correlation between the LBA parameter for non-decision time and the properties of the bilateral cortico-spinal tract.

To our eyes, these previous attempts are suboptimal, since mean, median or SD of RT are blunt summaries of a highly composite measure, while model estimates of non-decision time can be misguided, as we have shown here. We anticipate that the use of the approaches described in this article to isolate individual differences in 1) visual delay and 2) manual output time, will lead to more robust and specific patterns of correlation. In particular, using saccadic dip onset time would give us the best chance to assess the role of individual differences in optic radiation microstructure, while contrasting simple saccadic and manual reaction time distributions will do the same for the cortico-spinal tracts. Future studies will need to extend those approaches across the age span and clinical conditions, in order to form a better understanding of the contribution of these tracts to individual differences in response speed.

## Conclusion

We hope to have demonstrated that empirical approaches to estimating non-decision time are both straightforward and necessary for building solid ground for future sensorimotor decision research. In contrast to simple model fitting that can typically be applied to most RT datasets, the approaches described in this work generally require the acquisition of specific data. Future research will need to quantify key properties, such as test-retest repeatability as a function of trial numbers, which will in turn dictate the applicability of these methods to more challenging populations (e.g., children, patients) and inform the sample sizes required. The simplicity of the tasks proposed (simple RT or distractor task) makes them amenable to a large range of populations and, for human observers, do not require detailed instructions or training.

We hope this work will inspire a wide range of researchers and further our understanding of decision and its biological underpinning. For instance, researchers interested in understanding action decision through modelling may find these useful to assess the plausibility of models’ outcome and further constrain their model. Researchers interested in individual differences may want to directly use these measures, as alternatives or in addition, to model estimates when contrasting decision or non-decision time across individuals or between groups, or linking them to physiological measures such as genetics or brain structure. Further, researchers interested in unravelling the neural dynamics of these decisions may use these estimates to determine time windows of interest when analysing electrophysiological signals that possess high temporal resolution, such as single cell recordings, electroencephalography, magnetoencephalography or electromyograms, or to choose optimal stimulation times in brain simulation protocols such as transcranial magnetic or direct-current stimulation.

## CRediT Statement

**AB:** Lead on all aspects of the article, including conceptualisation, methodology, software, formal analysis, investigation, data curation, writing of original draft, review & editing, visualisation. **CH:** Conceptualisation, methodology, software, writing – review & editing. **PS:** Conceptualisation, funding acquisition, writing – review and editing.

## Acknowledgements

The authors acknowledge funding from the School of Psychology at Cardiff University, the Economic and Social Research Council (ES/K002325/1) and the Wellcome Trust (104943/Z/14/Z). Funding sources had no involvement at any stage of the reported research. We are grateful to Steven P. Errington, Jeff Schall, Antimo Buonocore and Alessandro Gulberti for sharing data, and to Georgie Powell for useful feedback on the manuscript. The original submission of this article was published on BIORXIV. Some of the results were presented at the Annual meeting of the Mathematical Psychology Association in 2020, the Christmas meeting of the Applied Vision Association in 2021 and the European Conference on Eye Movements in 2022. All analysis code used for the present article and all the data collected in Cardiff can be found on the OSF from [here](https://osf.io/gz9uc/?view_only=30c56b2c454949ddbe2b4d56da5f87f6).

## Footnotes

1 (“diffusion*model*” OR DDM OR “LBA model” OR “linear ballistic accumulator” OR “decision model*”)

2 Adding to the above: AND (“non-decision time” OR “non decision time” OR “nondecision time” OR “response execution time” OR “sensory time” OR “sensory delay” OR “sensory conduction delay” OR “sensory conduction time”)

3 https://github.com/smfleming/LBA

## Notes

https://osf.io/gz9uc/?view_only=30c56b2c454949ddbe2b4d56da5f87f6

### Competing Interest Statement

The authors have declared no competing interest.

## References

Aschenbrenner, A. J., Balota, D. A., Gordon, B. A., Ratcliff, R., & Morris, J. C. (2016). A diffusion model analysis of episodic recognition in preclinical individuals with a family history for Alzheimer’s disease: The adult children study. Neuropsychology, 30(2), 225–238. https://doi.org/10.1037/neu0000222

Boehm, U., Annis, J., Frank, M. J., Hawkins, G. E., Heathcote, A., Kellen, D., Krypotos, A.-M., Lerche, V., Logan, G. D., Palmeri, T. J., Van Ravenzwaaij, D., Servant, M., Singmann, H., Starns, J. J., Voss, A., Wiecki, T. V., Matzke, D., & Wagenmakers, E.-J. (2018). Estimating across-trial variability parameters of the Diffusion Decision Model: Expert advice and recommendations. Journal of Mathematical Psychology, 87, 46–75. https://doi.org/10.1016/j.jmp.2018.09.004

Bolz, J., Rosner, G., & Wassle, H. (1982). Response latency of brisk-sustained (X) and brisk-transient (Y) cells in the cat retina. J Physiol, 328, 171–190. https://doi.org/10.1113/jphysiol.1982.sp014258

Bompas, A., Campbell, A. E., & Sumner, P. (2020). Cognitive control and automatic interference in mind and brain: A unified model of saccadic inhibition and countermanding. Psychol Rev, 127(4), 524–561. https://doi.org/10.1037/rev0000181

Bompas, A., Hedge, C., & Sumner, P. (2017). Speeded saccadic and manual visuo-motor decisions: Distinct processes but same principles. Cogn Psychol, 94, 26–52. https://doi.org/10.1016/j.cogpsych.2017.02.002

Bompas, A., & Sumner, P. (2008). Sensory sluggishness dissociates saccadic, manual, and perceptual responses: An S-cone study. Journal of Vision, 8(8), 1–13. https://doi.org/10.1167/8.8.10

Bompas, A., & Sumner, P. (2009a). Oculomotor distraction by signals invisible to the retinotectal and magnocellular pathways. Journal of Neurophysiology, 102, 2387–2395. 10.1152/jn.00359.2009

Bompas, A., & Sumner, P. (2009b). Temporal dynamics of saccadic distraction. Journal of Vision, 9(9), 1–14. https://doi.org/10.1167/9.9.17

Bompas, A., & Sumner, P. (2011). Saccadic inhibition reveals the timing of automatic and voluntary signals in the human brain. Journal of Neuroscience, 31(35), 12501–12512. https://doi.org/10.1523/jneurosci.2234-11.2011

Booth, T., Dykiert, D., Corley, J., Gow, A. J., Morris, Z., Munoz Maniega, S., Royle, N. A., Del, C. V. H. M., Starr, J. M., Penke, L., Bastin, M. E., Wardlaw, J. M., & Deary, I. J. (2019). Reaction time variability and brain white matter integrity. Neuropsychology, 33(5), 642–657. https://doi.org/10.1037/neu0000483

Boucher, L., Palmeri, T. J., Logan, G. D., & Schall, J. D. (2007). Inhibitory control in mind and brain: An interactive race model of countermanding Saccades. Psychological Review, 114(2), 376–397. https://doi.org/10.1037/0033-295x.114.2.376

Boucher, L., Stuphorn, V., Logan, G. D., Schall, J. D., & Palmeri, T. J. (2007). Stopping eye and hand movements: Are the processes independent? Perception & Psychophysics, 69(5), 785–801. https://doi.org/10.3758/BF03193779

Brown, S. D., & Heathcote, A. (2008). The simplest complete model of choice response time: Linear ballistic accumulation. Cognitive Psychology, 57(3), 153–178. https://doi.org/10.1016/j.cogpsych.2007.12.002

Buonocore, A., & McIntosh, R. D. (2012). Modulation of saccadic inhibition by distractor size and location. Vision Research, 69, 32–41. https://doi.org/10.1016/J.Visres.2012.07.010

Buonocore, A., Tian, X., Khademi, F., & Hafed, Z. M. (2021). Instantaneous movement-unrelated midbrain activity modifies ongoing eye movements. Elife, 10. https://doi.org/10.7554/eLife.64150

Campbell, A. E., Chambers, C. D., Allen, C. P. G., Hedge, C., & Sumner, P. (2017). Impairment of manual but not saccadic response inhibition following acute alcohol intoxication. Drug Alcohol Depend, 181, 242–254. https://doi.org/10.1016/j.drugalcdep.2017.08.022

Carpenter, R. H. S., & Williams, M. L. L. (1995). Neural Computation of Log Likelihood in Control of Saccadic Eye-Movements. Nature, 377(6544), 59–62. https://doi.org/10.1038/377059a0

Cassey, P., Heathcote, A., & Brown, S. D. (2014). Brain and behavior in decision-making. PLoS Comput Biol, 10(7), e1003700. https://doi.org/10.1371/journal.pcbi.1003700

Chen, C. Y., Sonnenberg, L., Weller, S., Witschel, T., & Hafed, Z. M. (2018). Spatial frequency sensitivity in macaque midbrain. Nat Commun, 9(1), 2852. https://doi.org/10.1038/s41467-018-05302-5

Donkin, C., Brown, S., Heathcote, A., & Wagenmakers, E. J. (2011). Diffusion versus linear ballistic accumulation: different models but the same conclusions about psychological processes? Psychon Bull Rev, 18(1), 61–69. https://doi.org/10.3758/s13423-010-0022-4

Durst, M., & Janczyk, M. (2019). Two types of backward crosstalk: Sequential modulations and evidence from the diffusion model. Acta Psychol (Amst), 193, 132–152. https://doi.org/10.1016/j.actpsy.2018.11.013

Dutilh, G., Annis, J., Brown, S. D., Cassey, P., Evans, N. J., Grasman, R., Hawkins, G. E., Heathcote, A., Holmes, W. R., Krypotos, A. M., Kupitz, C. N., Leite, F. P., Lerche, V., Lin, Y. S., Logan, G. D., Palmeri, T. J., Starns, J. J., Trueblood, J. S., van Maanen, L., … Donkin, C. (2019). The Quality of Response Time Data Inference: A Blinded, Collaborative Assessment of the Validity of Cognitive Models. Psychon Bull Rev, 26(4), 1051–1069. https://doi.org/10.3758/s13423-017-1417-2

Dutilh, G., Krypotos, A. M., & Wagenmakers, E. J. (2011). Task-related versus stimulus-specific practice. Exp Psychol, 58(6), 434–442. https://doi.org/10.1027/1618-3169/a000111

Evans, N. J. (2021). Think fast! The implications of emphasizing urgency in decision-making. Cognition, 214, 104704. https://doi.org/10.1016/j.cognition.2021.104704

Evans, N. J., & Wagenmakers, E.-J. (2020). Evidence accumulation models: Current limitations and future directions. The Quantitative Methods for Psychology, 16, 73–90. https://doi.org/10.20982/tqmp.16.2.p073

Fjell, A. M., Westlye, L. T., Amlien, I. K., & Walhovd, K. B. (2011). Reduced white matter integrity is related to cognitive instability. J Neurosci, 31(49), 18060–18072. https://doi.org/10.1523/JNEUROSCI.4735-11.2011

Forstmann, B. U., Tittgemeyer, M., Wagenmakers, E. J., Derrfuss, J., Imperati, D., & Brown, S. (2011). The speed-accuracy tradeoff in the elderly brain: a structural model-based approach. J Neurosci, 31(47), 17242–17249. https://doi.org/10.1523/JNEUROSCI.0309-11.2011

Fosco, W. D., White, C. N., & Hawk, L. W., Jr. (2017). Acute Stimulant Treatment and Reinforcement Increase the Speed of Information Accumulation in Children with ADHD. J Abnorm Child Psychol, 45(5), 911–920. https://doi.org/10.1007/s10802-016-0222-0

Gold, J. I., & Shadlen, M. N. (2000). Representation of a perceptual decision in developing oculomotor commands. Nature, 404(6776), 390–394. https://doi.org/10.1038/35006062

Gomez, P., Ratcliff, R., & Childers, R. (2015). Pointing, looking at, and pressing keys: A diffusion model account of response modality. J Exp Psychol Hum Percept Perform, 41(6), 1515–1523. https://doi.org/10.1037/a0039653

Grange, J. A., & Schuch, S. (2021). A spurious correlation between difference scores in evidence accumulation model parameters. PsycArXiv. https://doi.org/https://doi.org/10.31234/osf.io/u6py8

Gulberti, A., Arndt, P. A., & Colonius, H. (2014). Stopping eyes and hands: evidence for non-independence of stop and go processes and for a separation of central and peripheral inhibition. Front Hum Neurosci, 8, 61. https://doi.org/10.3389/fnhum.2014.00061

Haasz, J., Westlye, E. T., Fjaer, S., Espeseth, T., Lundervold, A., & Lundervold, A. J. (2013). General fluid-type intelligence is related to indices of white matter structure in middle-aged and old adults. Neuroimage, 83, 372–383. https://doi.org/10.1016/j.neuroimage.2013.06.040

Hedge, C., Powell, G., Bompas, A., & Sumner, P. (2020). Self-reported impulsivity does not predict response caution. Pers Individ Dif, 167, 110257. https://doi.org/10.1016/j.paid.2020.110257

Hedge, C., Powell, G., Bompas, A., & Sumner, P. (2022). Strategy and processing speed eclipse individual differences in control ability in conflict tasks. J Exp Psychol Learn Mem Cogn, 48(10), 1448–1469. https://doi.org/10.1037/xlm0001028

Hedge, C., Powell, G., & Sumner, P. (2018). The reliability paradox: Why robust cognitive tasks do not produce reliable individual differences. Behav Res Methods, 50(3), 1166–1186. https://doi.org/10.3758/s13428-017-0935-1

Heitz, R. P., & Schall, J. D. (2012). Neural mechanisms of speed-accuracy tradeoff. Neuron, 76(3), 616–628. https://doi.org/10.1016/j.neuron.2012.08.030

Huang, Y. T., Georgiev, D., Foltynie, T., Limousin, P., Speekenbrink, M., & Jahanshahi, M. (2015). Different effects of dopaminergic medication on perceptual decision-making in Parkinson’s disease as a function of task difficulty and speed-accuracy instructions. Neuropsychologia, 75, 577–587. https://doi.org/10.1016/j.neuropsychologia.2015.07.012

Imburgio, M. J., & Orr, J. M. (2021). Component processes underlying voluntary task selection: Separable contributions of task-set inertia and reconfiguration. Cognition, 212, 104685. https://doi.org/10.1016/j.cognition.2021.104685

Imms, P., Dominguez, D. J., Burmester, A., Seguin, C., Clemente, A., Dhollander, T., Wilson, P. H., Poudel, G., & Caeyenberghs, K. (2021). Navigating the link between processing speed and network communication in the human brain. Brain Struct Funct, 226(4), 1281–1302. https://doi.org/10.1007/s00429-021-02241-8

Karahan, E., Costigan, A. G., Graham, K. S., Lawrence, A. D., & Zhang, J. (2019). Cognitive and White-Matter Compartment Models Reveal Selective Relations between Corticospinal Tract Microstructure and Simple Reaction Time. J Neurosci, 39(30), 5910–5921. https://doi.org/10.1523/jneurosci.2954-18.2019

Karalunas, S. L., Geurts, H. M., Konrad, K., Bender, S., & Nigg, J. T. (2014). Annual research review: Reaction time variability in ADHD and autism spectrum disorders: measurement and mechanisms of a proposed trans-diagnostic phenotype. J Child Psychol Psychiatry, 55(6), 685–710. https://doi.org/10.1111/jcpp.12217

Karalunas, S. L., Hawkey, E., Gustafsson, H., Miller, M., Langhorst, M., Cordova, M., Fair, D., & Nigg, J. T. (2018). Overlapping and Distinct Cognitive Impairments in Attention-Deficit/Hyperactivity and Autism Spectrum Disorder without Intellectual Disability. J Abnorm Child Psychol, 46(8), 1705–1716. https://doi.org/10.1007/s10802-017-0394-2

Kelly, S. P., Corbett, E. A., & O’Connell, R. G. (2021). Neurocomputational mechanisms of prior-informed perceptual decision-making in humans. Nat Hum Behav, 5(4), 467–481. https://doi.org/10.1038/s41562-020-00967-9

Kievit, R. A., Davis, S. W., Griffiths, J., Correia, M. M., Cam, C., & Henson, R. N. (2016). A watershed model of individual differences in fluid intelligence. Neuropsychologia, 91, 186–198. https://doi.org/10.1016/j.neuropsychologia.2016.08.008

Kuznetsova, K. A., Maniega, S. M., Ritchie, S. J., Cox, S. R., Storkey, A. J., Starr, J. M., Wardlaw, J. M., Deary, I. J., & Bastin, M. E. (2016). Brain white matter structure and information processing speed in healthy older age. Brain Struct Funct, 221(6), 3223–3235. https://doi.org/10.1007/s00429-015-1097-5

Lee, J., Kim, H. R., & Lee, C. (2010). Trial-to-trial variability of spike response of V1 and saccadic response time. J Neurophysiol, 104(5), 2556–2572. https://doi.org/jn.01040.2009

Lerche, V., & Voss, A. (2016). Model Complexity in Diffusion Modeling: Benefits of Making the Model More Parsimonious. Front Psychol, 7, 1324. https://doi.org/10.3389/fpsyg.2016.01324

Lerche, V., & Voss, A. (2017). Retest reliability of the parameters of the Ratcliff diffusion model. Psychol Res, 81(3), 629–652. https://doi.org/10.1007/s00426-016-0770-5

Lerche, V., & Voss, A. (2018). Speed-accuracy manipulations and diffusion modeling: Lack of discriminant validity of the manipulation or of the parameter estimates? Behav Res Methods, 50(6), 2568–2585. https://doi.org/10.3758/s13428-018-1034-7

Li, X., & Basso, M. A. (2008). Preparing to move increases the sensitivity of superior colliculus neurons. J Neurosci, 28(17), 4561–4577. https://doi.org/10.1523/jneurosci.5683-07.2008

Lo, C. C., Boucher, L., Pare, M., Schall, J. D., & Wang, X. J. (2009). Proactive Inhibitory Control and Attractor Dynamics in Countermanding Action: A Spiking Neural Circuit Model. Journal of Neuroscience, 29(28), 9059–9071. https://doi.org/10.1523/Jneurosci.6164-08.2009

Marino, R. A., Levy, R., Boehnke, S., White, B. J., Itti, L., & Munoz, D. P. (2012). Linking visual response properties in the superior colliculus to saccade behavior. Eur J Neurosci, 35(11), 1738–1752. https://doi.org/10.1111/j.1460-9568.2012.08079.x

Merkt, J., Singmann, H., Bodenburg, S., Goossens-Merkt, H., Kappes, A., Wendt, M., & Gawrilow, C. (2013). Flanker performance in female college students with ADHD: a diffusion model analysis. Atten Defic Hyperact Disord, 5(4), 321–341. https://doi.org/10.1007/s12402-013-0110-1

Miletić, S., Turner, B. M., Forstmann, B. U., & van Maanen, L. (2017). Parameter recovery for the leaky competing accumulator model. Journal of Mathematical Psychology, 76, 25–50. https://doi.org/10.1016/j.jmp.2016.12.001

Miyashita, N., & Hikosaka, O. (1996). Minimal synaptic delay in the saccadic output pathway of the superior colliculus studied in awake monkey. Exp Brain Res, 112(2), 187–196. https://doi.org/10.1007/BF00227637

Munoz, D. P., & Wurtz, R. H. (1993). Fixation cells in monkey superior colliculus. II. Reversible activation and deactivation. J Neurophysiol, 70(2), 576–589. https://doi.org/10.1152/jn.1993.70.2.576

Patanaik, A., Zagorodnov, V., Kwoh, C. K., & Chee, M. W. (2014). Predicting vulnerability to sleep deprivation using diffusion model parameters. Journal of Sleep Research, 23(5), 576–584. https://doi.org/10.1111/jsr.12166

Penke, L., Maniega, S. M., Bastin, M. E., Valdes Hernandez, M. C., Murray, C., Royle, N. A., Starr, J. M., Wardlaw, J. M., & Deary, I. J. (2012). Brain white matter tract integrity as a neural foundation for general intelligence. Mol Psychiatry, 17(10), 1026–1030. https://doi.org/10.1038/mp.2012.66

Penke, L., Munoz Maniega, S., Murray, C., Gow, A. J., Hernandez, M. C., Clayden, J. D., Starr, J. M., Wardlaw, J. M., Bastin, M. E., & Deary, I. J. (2010). A general factor of brain white matter integrity predicts information processing speed in healthy older people. J Neurosci, 30(22), 7569–7574. https://doi.org/10.1523/jneurosci.1553-10.2010

Powell, G., Jones, C. R. G., Hedge, C., Charman, T., Happe, F., Simonoff, E., & Sumner, P. (2019). Face processing in autism spectrum disorder re-evaluated through diffusion models. Neuropsychology, 33(4), 445–461. https://doi.org/10.1037/neu0000524

Purcell, B. A., Heitz, R. P., Cohen, J. Y., Schall, J. D., Logan, G. D., & Palmeri, T. J. (2010). Neurally Constrained Modeling of Perceptual Decision Making. Psychological Review, 117(4), 1113–1143. https://doi.org/10.1037/A0020311

Ratcliff, R. (2013). Parameter variability and distributional assumptions in the diffusion model. Psychol Rev, 120(1), 281–292. https://doi.org/10.1037/a0030775

Ratcliff, R., & Rouder, J. N. (1998). Modeling response times for two-choice decisions. Psychological Science, 9(5), 347–356. https://doi.org/10.1111/1467-9280.00067

Ratcliff, R., & Tuerlinckx, F. (2002). Estimating parameters of the diffusion model: approaches to dealing with contaminant reaction times and parameter variability. Psychon Bull Rev, 9(3), 438–481. https://doi.org/10.3758/bf03196302

Ray, S., Pouget, P., & Schall, J. D. (2009). Functional Distinction Between Visuomovement and Movement Neurons in Macaque Frontal Eye Field During Saccade Countermanding. Journal of Neurophysiology, 102(6), 3091–3100. https://doi.org/10.1152/Jn.00270.2009

Reingold, E. M., & Stampe, D. M. (2002). Saccadic inhibition in voluntary and reflexive saccades. J Cogn Neurosci, 14(3), 371–388. https://doi.org/10.1162/089892902317361903

Reingold, E. M., & Stampe, D. M. (2004). Saccadic inhibition in reading. Journal of Experimental Psychology-Human Perception and Performance, 30(1), 194–211. https://doi.org/10.1037/0096-1523.30.1.194

Reppert, T. R., Servant, M., Heitz, R. P., & Schall, J. D. (2018). Neural mechanisms of speed-accuracy tradeoff of visual search: saccade vigor, the origin of targeting errors, and comparison of the superior colliculus and frontal eye field. J Neurophysiol, 120(1), 372–384. https://doi.org/10.1152/jn.00887.2017

Rohner, J., & Lai, C. K. (2021). A Diffusion Model Approach for Understanding the Impact of 17 Interventions on the Race Implicit Association Test. Pers Soc Psychol Bull, 47(9), 1374–1389. https://doi.org/10.1177/0146167220974489

Sajad, A., Godlove, D. C., & Schall, J. D. (2019). Cortical microcircuitry of performance monitoring. Nat Neurosci, 22(2), 265–274. https://doi.org/10.1038/s41593-018-0309-8

Salinas, E., & Stanford, T. R. (2018). Saccadic inhibition interrupts ongoing oculomotor activity to enable the rapid deployment of alternate movement plans. Sci Rep, 8(1), 14163. https://doi.org/10.1038/s41598-018-32224-5

Salinas, E., & Stanford, T. R. (2021). Under time pressure, the exogenous modulation of saccade plans is ubiquitous, intricate, and lawful. Curr Opin Neurobiol, 70, 154–162. https://doi.org/10.1016/j.conb.2021.10.012

Sandry, J., & Ricker, T. J. (2022). Motor speed does not impact the drift rate: a computational HDDM approach to differentiate cognitive and motor speed. Cogn Res Princ Implic, 7(1), 66. https://doi.org/10.1186/s41235-022-00412-7

Servant, M., Logan, G. D., Gajdos, T., & Evans, N. J. (2021). An integrated theory of deciding and acting. J Exp Psychol Gen, 150(12), 2435–2454. https://doi.org/10.1037/xge0001063

Servant, M., Tillman, G., Schall, J. D., Logan, G. D., & Palmeri, T. J. (2019). Neurally constrained modeling of speed-accuracy tradeoff during visual search: gated accumulation of modulated evidence. J Neurophysiol, 121(4), 1300–1314. https://doi.org/10.1152/jn.00507.2018

Servant, M., White, C., Montagnini, A., & Burle, B. (2016). Linking Theoretical Decision-making Mechanisms in the Simon Task with Electrophysiological Data: A Model-based Neuroscience Study in Humans. J Cogn Neurosci, 28(10), 1501–1521. https://doi.org/10.1162/jocn_a_00989

Smith, P. L. (1990). A note on the distribution of response times for a random walk with Gaussian increments. Journal of Mathematical Psychology, 34, 445–459.

Stampe, D. M., & Reingold, E. M. (2002). Influence of stimulus characteristics on the latency of saccadic inhibition. Prog Brain Res, 140, 73–87. https://doi.org/10.1016/S0079-6123(02)40043-X

Tamnes, C. K., Fjell, A. M., Westlye, L. T., Ostby, Y., & Walhovd, K. B. (2012). Becoming consistent: developmental reductions in intraindividual variability in reaction time are related to white matter integrity. J Neurosci, 32(3), 972–982. https://doi.org/10.1523/jneurosc.4779-11.2012

Team, J. (2020). JASP (Version 0.14.1).

Teodorescu, A. R., & Usher, M. (2013). Disentangling decision models: from independence to competition. Psychol Rev, 120(1), 1–38. https://doi.org/10.1037/a0030776

Theisen, M., Lerche, V., von Krause, M., & Voss, A. (2021). Age differences in diffusion model parameters: a meta-analysis. Psychol Res, 85(5), 2012–2021. https://doi.org/10.1007/s00426-020-01371-8

Trappenberg, T. P., Dorris, M. C., Munoz, D. P., & Klein, R. M. (2001). A model of saccade initiation based on the competitive integration of exogenous and endogenous signals in the superior colliculus. Journal of Cognitive Neuroscience, 13(2), 256–271. https://doi.org/10.1162/089892901564306

Ulrichsen, K. M., Alnaes, D., Kolskar, K. K., Richard, G., Sanders, A. M., Dorum, E. S., Ihle-Hansen, H., Pedersen, M. L., Tornas, S., Nordvik, J. E., & Westlye, L. T. (2020). Dissecting the cognitive phenotype of post-stroke fatigue using computerized assessment and computational modeling of sustained attention. Eur J Neurosci, 52(7), 3828–3845. https://doi.org/10.1111/ejn.14861

van Ravenzwaaij, D., Donkin, C., & Vandekerckhove, J. (2017). The EZ diffusion model provides a powerful test of simple empirical effects. Psychon Bull Rev, 24(2), 547–556. https://doi.org/10.3758/s13423-016-1081-y

van Ravenzwaaij, D., Dutilh, G., & Wagenmakers, E. J. (2012). A diffusion model decomposition of the effects of alcohol on perceptual decision making. Psychopharmacology (Berl), 219(4), 1017–1025. https://doi.org/10.1007/s00213-011-2435-9

van Ravenzwaaij, D., & Oberauer, K. (2009). How to use the diffusion model: Parameter recovery of three methods: Ez, fast-dm, and DMAT. Journal of Mathematical Psychology, 53(6), 463–473. https://doi.org/10.1016/j.jmp.2009.09.004

Van Zandt, T., & Ratcliff, R. (1995). Statistical mimicking of reaction time data: Single-process models, parameter variability, and mixtures. Psychon Bull Rev, 2(1), 20–54. https://doi.org/10.3758/BF03214411

Vandekerckhove, J., & Tuerlinckx, F. (2008). Diffusion model analysis with MATLAB: a DMAT primer. Behav Res Methods, 40(1), 61–72. https://doi.org/10.3758/brm.40.1.61

Verdonck, S., & Tuerlinckx, F. (2016). Factoring out nondecision time in choice reaction time data: Theory and implications. Psychol Rev, 123(2), 208–218. https://doi.org/10.1037/rev0000019

Voss, A., Rothermund, K., & Voss, J. (2004). Interpreting the parameters of the diffusion model: an empirical validation. Mem Cognit, 32(7), 1206–1220. https://doi.org/10.3758/bf03196893

Wagenmakers, E. J., van der Maas, H. L., & Grasman, R. P. (2007). An EZ-diffusion model for response time and accuracy. Psychon Bull Rev, 14(1), 3–22. https://doi.org/10.3758/bf03194023

Wagner, B., Clos, M., Sommer, T., & Peters, J. (2020). Dopaminergic Modulation of Human Intertemporal Choice: A Diffusion Model Analysis Using the D2-Receptor Antagonist Haloperidol. J Neurosci, 40(41), 7936–7948. https://doi.org/10.1523/jneurosci.0592-20.2020

Weindel, G., Gajdos, T., Burle, B., & Alario, F. X. (2021). The Decisive Role of Non-Decision Time for Interpreting the Parameters of Decision Making Models. PsyArXiv. https://doi.org/ https://doi.org/10.31234/osf.io/gewb3

White, B. J., Boehnke, S. E., Marino, R. A., Itti, L., & Munoz, D. P. (2009). Color-Related Signals in the Primate Superior Colliculus. Journal of Neuroscience, 29(39), 12159–12166. https://doi.org/10.1523/Jneurosci.1986-09.2009

White, C. N., Servant, M., & Logan, G. D. (2018). Testing the validity of conflict drift-diffusion models for use in estimating cognitive processes: A parameter-recovery study. Psychon Bull Rev, 25(1), 286–301. https://doi.org/10.3758/s13423-017-1271-2

Yang, Y., Bender, A. R., & Raz, N. (2015). Age related differences in reaction time components and diffusion properties of normal-appearing white matter in healthy adults. Neuropsychologia, 66, 246–258. https://doi.org/10.1016/j.neuropsychologia.2014.11.020

Zhang, J., Rittman, T., Nombela, C., Fois, A., Coyle-Gilchrist, I., Barker, R. A., Hughes, L. E., & Rowe, J. B. (2016). Different decision deficits impair response inhibition in progressive supranuclear palsy and Parkinson’s disease. Brain, 139(Pt 1), 161–173. https://doi.org/10.1093/brain/awv331

Ziv, I., & Bonneh, Y. S. (2021). Oculomotor inhibition during smooth pursuit and its dependence on contrast sensitivity. J Vis, 21(2), 12. https://doi.org/10.1167/jov.21.2.12

